# *Garamaudo bauciensis*, a new freshwater Mosasauridae (Reptilia, Squamata) from the Santonian (Late Cretaceous) of Provence, southeastern France

**DOI:** 10.64898/2025.12.11.693649

**Authors:** Nathalie Bardet, Alexandra Houssaye, François-Louis Pelissier, Yves Dutour, Éric Turini, Thierry Tortosa

**Affiliations:** CR2P, Centre de Recherche en Paléontologie – Paris, UMR 7207 CNRS/MNHN/SU, Muséum National d’Histoire Naturelle – Paris, France; MECADEV, Mécanismes adaptatifs et Evolution, UMR 7179 CNRS/MNHN, Muséum National d’Histoire Naturelle – Paris, France; Muséum départemental du Var – Toulon, France; Muséum d’Histoire Naturelle – Aix-en-Provence, France; Réserve Naturelle Nationale Sainte-Victoire, Direction de l’Environnement, des Grands Projets et de la Recherche du Département des Bouches-du-Rhône – Marseille, France

**Keywords:** Mosasauroidea, Tethysaurinae, new taxon, Late Cretaceous, freshwater environment, southeastern France

## Abstract

Mosasauroids are primarily known as aquatic squamates that diversified in the marine realm during the Late Cretaceous. While most species had paddles, an elongated skull and body, and a laterally compressed tail, and lived in the open sea, some early species, with limbs similar to those of terrestrial animals and a varanoid-like body, lived in marine coastal shallow waters. Recently, mosasauroid remains have been discovered in freshwater environments in several continental deposits of the Santonian-Campanian of Europe. This study describes new cranial and postcranial mosasauroid remains from Santonian continental deposits in Provence, southeastern France, and attributes them to a new tethysaurine: *Garamaudo bauciensis* gen. et sp. nov. This mosasauroid, approximately 2.5 m long, with limbs looking adapted for a terrestrial locomotion and a sacrum, and with osteosclerosis in its vertebrae and humerus (at least), probably hovered at shallow depth in freshwater ecosystems. This new discovery reveals that it is only within the Tethysaurinae that multiple freshwater forms evolved, pointing to possible niche partitioning, associated with the retention of the plesiopedal / plesiopelvic condition, in parallel with the invasion of open waters by most other hydropedal and hydropelvic mosasauroids. These considerations suggest a richer evolutionary history of mosasauroids than is currently assumed.

## Introduction

Mosasauroidea is an extinct group of aquatic squamates that invaded and diversified in the aquatic environment at the beginning of the Cenomanian, spreading worldwide from the Santonian to the end of the Maastrichtian, and becoming extinct during the biological crisis of the K/Pg event (e.g., Bardet *et al*., 2014; Polcyn *et al*., 2014). They are usually regarded as toxicoferans (Toxicofera including Anguimorpha, Iguania, Serpentes and all taxa closer to these taxa than to other squamates), and either as the sister group of Varanoidea (Polcyn *et al*., 2022) or Serpentes (Simoes *et al*., 2021). Mosasauroidea possess elongated skulls and bodies, and long, laterally compressed tails. They encompass a high diversity of sizes, from 1 to 15 m in length, in ecology, from freshwater, shallow marine to open waters, and in diet, from mollusks and arthropods to large fish and reptiles (e.g., Bardet *et al*., 2025).

Most mosasauroids possess a morphology strongly modified for an aquatic lifestyle, with paddles and the loss of connection between the vertebral column and the pelvis—that is, the absence of a true sacrum. This corresponds to the so-called hydropedal and hydropelvic grade (Caldwell & Palci, 2007). Conversely, more primitive forms display limbs looking adapted for a terrestrial locomotion and a sacrum, illustrating a plesiopedal and plesiopelvic condition. An intermediary pattern with plesiopedal limbs but no sacrum illustrates a third ecological grade (Bell & Polcyn, 2005; Caldwell & Palci, 2007).

Most mosasauroid remains are from marine deposits. However, a few taxa and undetermined mosasauroid remains have been encountered in freshwater environments, such as shallow estuarine and fluvial deposits (Holmes *et al*., 1999). Particularly common are freshwater occurrences in the Santonian-Campanian continental deposits of Europe, from Portugal to Hungary and Austria, passing through southern France (Makádi *et al*., 2012; Garcia *et al*., 2015; Ősi *et al*., 2019, 2021).

This study focuses on new mosasauroid remains discovered in Santonian continental deposits of Provence, southeastern France. These deposits are geographically and stratigraphically close to those lower Campanian deposits in Languedoc, southern France, where the Villeveyrac mosasauroid was discovered (Garcia *et al*., 2015). This work presents a description of this new material and its phylogenetic analysis, in order to determine its taxonomical identity and phylogenetic position within Mosasauroidea. We also describe the geological context and the microanatomy of some of the bones, in order to assess the paleobiology and paleoecology of the new taxon.

## Geological and Paleoenvironmental contexts

The Bouc-Bel-Air - Sousquières (BBAS) locality (Figure 1) is situated along departmental road RD8, in the city of Bouc-Bel-Air (Bouches-du-Rhône Department, Provence, southeastern France). It lies in the central part of the Aix-en-Provence sedimentary Basin, a broad east–west-oriented syncline formed in the context of Pyrenean-Provençal subsidence during the Late Cretaceous (Guieu, 1968; Gaviglio, 1985; Tortosa & Leleu, 2021).

**Figure 1.**
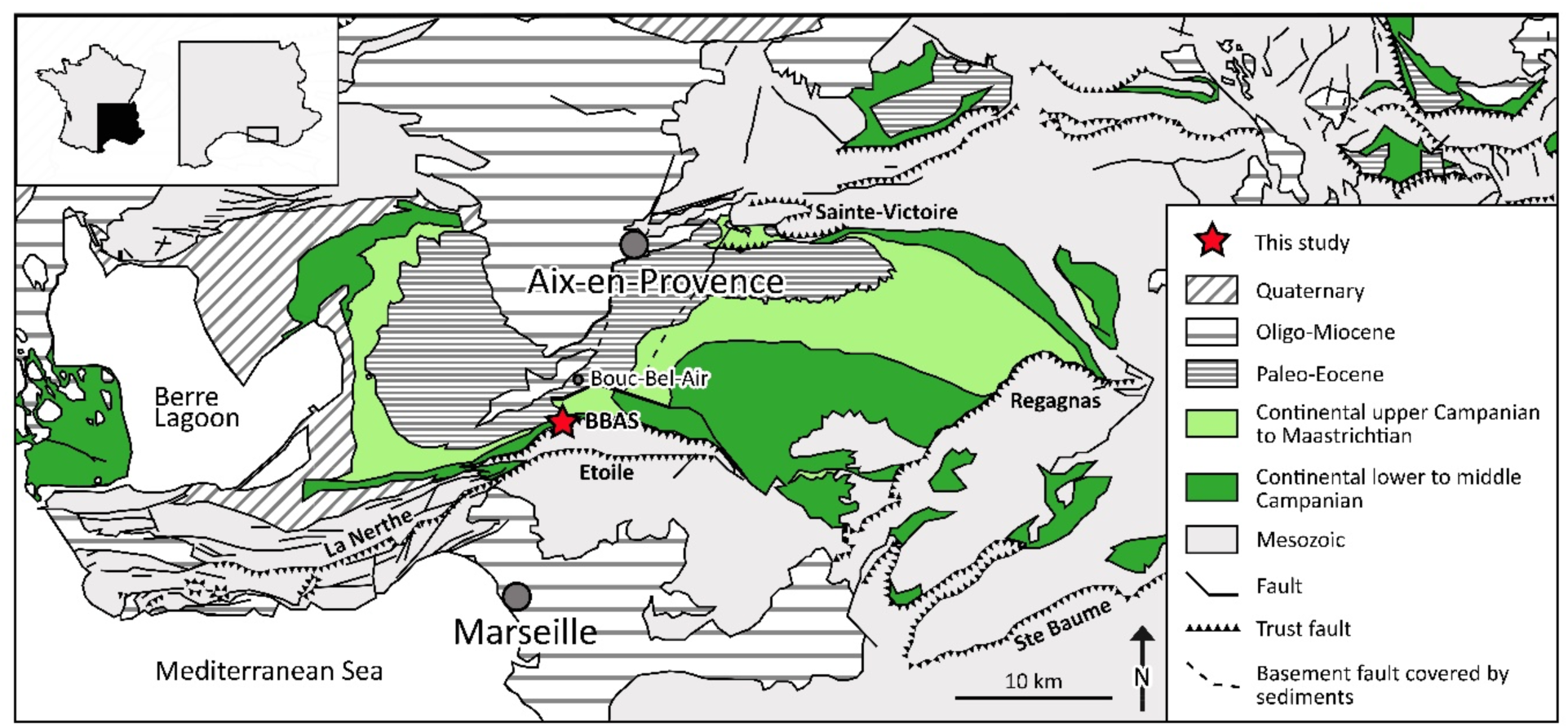
Geographical location of the Bouc-Bel-Air - Sousquières (BBAS) outcrop in the framework of the Upper Cretaceous continental deposits of the Aix-en-Provence Basin, Provence, southeastern France (after Tortosa *et al*., 2014).

The formations in this area form a fluvio-lacustrine complex deposited under a compressive Campanian setting (Gaviglio, 1985; Guieu, 1968). Two basal lithostratigraphic units compose the continental Campanian–Maastrichtian succession: the Valdonnian and the Fuvelian, both historically regarded as local stratigraphic stages (Figure 2C; Matheron, 1878; Collot, 1890; Babinot & Durand, 1980). This area, located along the so-called “Plan-de-Campagne – Bouc-Bel-Air flexure” has only been studied in an unpublished work by Honoré (1938), who described a reference section in the La Malle sector (∼2.8 km to the southwest).

**Figure 2.**
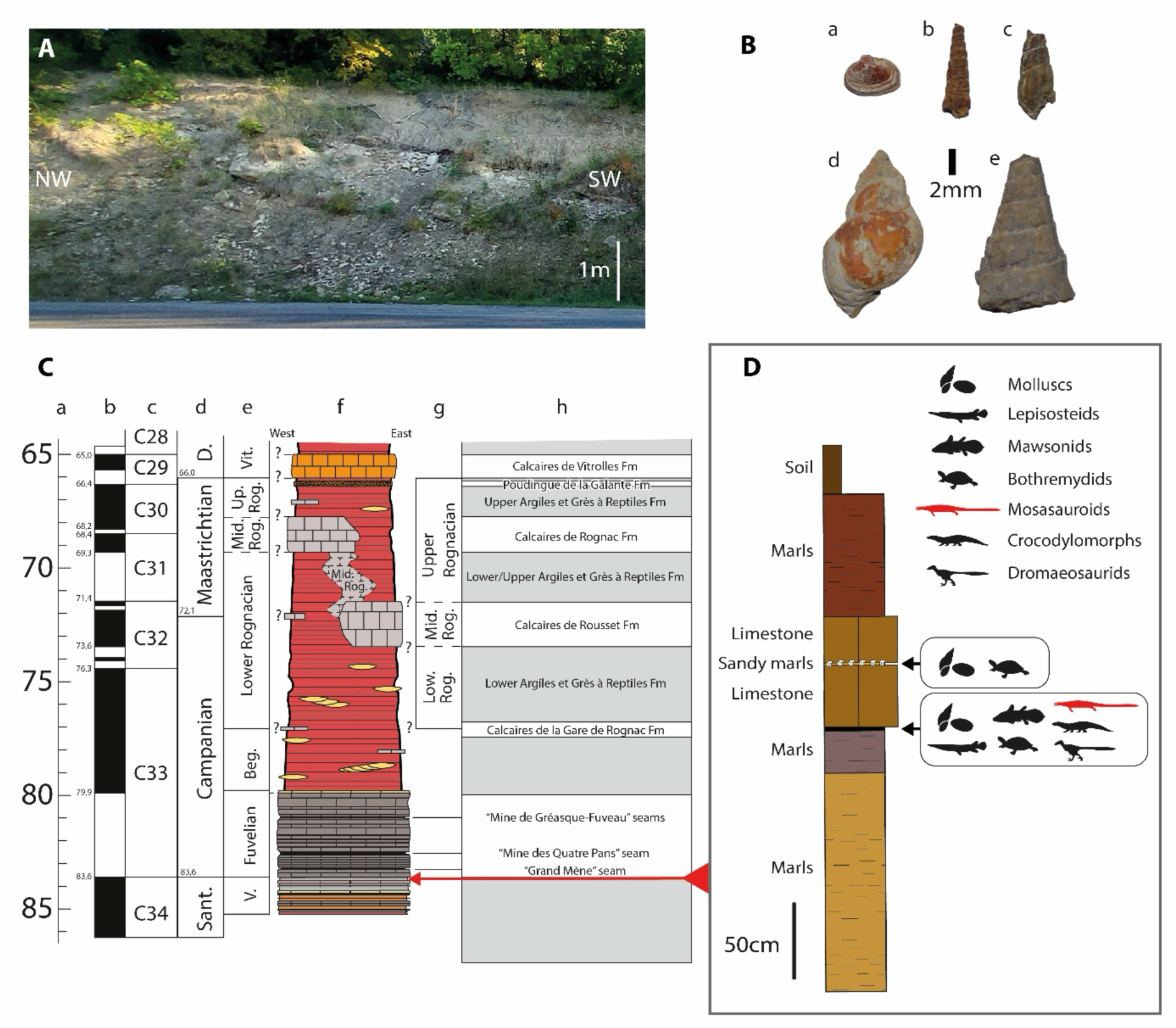
Stratigraphical details of the Bouc-Bel-Air - Sousquières (BBAS) outcrop. **A**, photograph of the site where *Garamaudo* gen. nov. was unearthed. **B**, molluscan assemblage found at BBSA, with *Corbicula concinna* (B-a), *Hadraxon acicula* (B-b), “*Melania vasseuri*” (B-c)*, Viviparus bosquiana* (B-d) and *Melania colloti* (B-e). **C**, stratigraphical position of BBAS in time scale with standard ages (C-a), geomagnetic polarity (C-b), chronozones (C-c), marine stratigraphy (C-d) and local continental facies (C-e, C-g), synthetic log of the Upper Cretaceous continental deposits of Aix-en-Provence Basin (C-f) with main continental formations (C-h), modified from Cojan & Moreau (2006), Gradstein *et al*. (2020) and Tortosa & Leleu (2021). **D**, detailed log and fossil-bearing horizons.

The BBAS outcrop is rather limited in size, characterized by alternations of beige to brown marls and grey lacustrine limestone beds (Figure 2A, D). Fossils are concentrated at the base of the limestones, at the contact with the marls, which are rich in organic matter and lacustrine mollusk shells, typical of the Valdonnian–Fuvelian assemblages (Figure 2B). These mollusks include the bivalve *Corbicula concinna* and the gastropods *Hadraxon acicula* and *Viviparus bosquiana* (Matheron, 1842; Honoré, 1938; Fabre-Taxy, 1951). To these species can be added *Melania colloti* (Roule, 1886; Fabre-Taxy, 1951), only known from the Valdonnian. Another unpublished form is recognized under the name “*Melania vasseuri*” (Honoré, 1938) and was previously misidentified as *M. praelonga* in Cavin *et al*. (2020). This latter taxon is so far known only from the levels of La Malle and Sousquières (TT., pers. obs.).

The associated vertebrate assemblage includes the actinopterygian *Atractosteus* cf. *africanus* (Cavin *et al*., 1996), the mawsoniid coelacanth *Axelrodichthys megadromos* (Cavin *et al*., 2020), a mosasauroid (this study), the bothremydid turtle cf. *Polysternon* sp., indeterminate crocodylomorphs and the dromaeosaurid cf. *Richardoestesia gilmorei* (TT., pers. obs.).

Direct correlations with the well-documented Valdonnian–Fuvelian succession of the Gardanne coal Basin (Caillol *et al*., 1988; Gonzalez, 1990) remain difficult because of the local tectonic context. The site lies on a structural slice bounded to the north by the Gardanne unit (limited by the Safre Fault) and to the south by the Simiane–Mimet and Sousquières thrust sheets (Guieu, 1968). The section described by Honoré (1938) at La Malle is lithologically similar to that of BBAS, and their fossil assemblages are directly comparable (Honoré, 1938; Fabre-Taxy, 1951). The absence of massive Fuvelian limestones in the immediate surroundings suggests either a rapid lateral facies variation toward more marly deposits, or a slightly older stratigraphic position corresponding to the transition between the Valdonnian and Fuvelian units (Dufaure *et al*., 1967; Babinot & Durand, 1980). The presence of *Melania colloti*, together with the tectonic compressions observed along the southern margin of the Gardanne Basin, support the latter hypothesis (Durand & Guieu, 1980). Pending further geological investigations, the BBAS locality is here interpreted as lying within the transition between the upper marly Valdonnian facies and the lower calcareous Fuvelian facies, in agreement with previous preliminary work on this locality (Cavin *et al*., 2020). This interval corresponds to a major episode of continental carbonate sedimentation, coeval with the progressive emergence of Provence and the establishment of an extensive endorheic lacustrine system in the Aix-en-Provence basin (Durand & Guieu, 1980; Tortosa & Leleu, 2021).

In details, the molluscan assemblage closely mirrors classical Valdonnian–basal Fuvelian faunas from Provence (Fabre-Taxy, 1951), dominated by *Melania*-type thiarids and *Hadraxon*, both of which indicate shallow, low-energy lacustrine to palustrine environments with organic-rich muds that favor the preservation of articulated vertebrate skeletons (Fabre-Taxy, 1951; Babinot & Durand, 1980; Cavin *et al*., 2020; this study). *Viviparus bosquiana* and *Cyanocyclas cuneata* typically occur along vegetated lake margins, implying clear, bicarbonate-rich waters, whereas *Corbicula* species inhabit quiet, sometimes poorly oxygenated floodplain ponds with episodic eutrophic conditions, consistent with shell concentrations at marl–limestone interfaces (Fabre-Taxy, 1951). Altogether, this assemblage points to a moderately eutrophic, carbonate-producing freshwater lake periodically influenced by flooding and shallowing events. This setting contrasts with the marshier, lignite-rich upper Fuvelian faunas of the Gardanne coal Basin, reflecting more humid, swampy environments (Fabre-Taxy, 1951; Babinot & Durand, 1980; Tortosa & Leleu, 2021).

Magnetostratigraphic studies suggest a Santonian age for the site (Westphal & Durand, 1990; corrected from Cojan & Moreau, 2006) (Figure 2C), making it one of the oldest Upper Cretaceous continental vertebrate localities of southeastern France, along with Ventabren – Air de Repos (Cavin *et al*., 2020) and Belcodène – Autoroute Diffuseur (Tortosa, 2014).

## Material and methods

### Material

The material consists of at least four individuals. The first consists of about 20 bones (CD13-PAL.2018.3.1.1 to 3, CD13-PAL.2018.3.1.5 to 15 and CD13-PAL.2018.3.1.18), including cranial (posterior mandibular unit), axial (cervical vertebrae and ribs) and appendicular (pelvis) elements. The sizes of the bones, the absence of duplicates, and the fact that they have been found in close association during one excavation campaign, undertaken by the Muséum d’Histoire Naturelle of Aix-en-Provence (MHNA) in 1997, are consistent with them being derived from a single individual. These remains are considered here as the holotype of *Garamaudo bauciencis*.

Two bones – the dorsal vertebra CD13-PAL.2018.3.1.4 and the humerus CD13-PAL.2018.3.1.16 – although found together with the holotype, are respectively much larger and smaller than the holotype, suggesting that they belong to two additional individuals.

Finally, the dorsal vertebra CD13-PAL.2018.3.4 is comparable in size to the holotype, but was found during a rescue campaign two decades later (2018), about 20 m from the *locus typicus*, and thus possibly represents an additional individual.

The microanatomical features of the various bones analyzed are consistent, whatever their size and occurrence, supporting the hypothesis that all bones probably belong to the same taxon, although to different individuals.

As such, the two dorsal vertebrae are considered referred specimens, and the humerus, whose underived features are consistent with the plesiomorphic morphology of the holotype bones (especially of the posterior mandibular unit and pelvis), is considered the paratype of *Garamaudo* gen. nov.

Some vertebral fragments (CD13-PAL.2018.3.1.17, CD13-PAL.2018.3.2, CD13-PAL.2018.3.3), unearthed in the *locus typicus*, are too incomplete to be confidently referred to the new taxon.

### Comparative Anatomy

Bone comparisons were made using the existing literature for extinct squamates (mosasauroids, ophidiomorphs) and extant *Varanus* skeletons from the MNHN Comparative Anatomy Collections (MNHN AC 1910-12, MNHN AC 1983-6).

The original positions of the vertebrae along the vertebral column were determined based on comparisons with modern varanids and fossil pythonomorphs Cope, 1869 (*sensu* Lee, 1997).

### CT-scanning

Three vertebrae (CD13-PAL.2018.3.1.1 and CD13-PAL.2018.3.1.3, holotype; CD13-PAL.2018.3.4, referred specimen), one rib (CD13-PAL.2018.3.1.13, holotype) and the humerus (CD13-PAL.2018.3.1.16, paratype) were scanned using high-resolution computed tomography at the MRI platform, hosted at the ISEM, University of Montpellier (UMR 5554; EasyTom 40-150, RX Solutions), with reconstructions performed using X-Act (RX Solutions). Voxel size naturally varies pending on specimen size: 17.8 µm (humerus and vertebra CD13-PAL.2018.3.4), 23.1 µm (vertebra CD13-PAL.2018.3.1.1), 21.5 µm (vertebra CD13-PAL.2018.3.1.3) and 24.2 µm (rib).

Image segmentation and visualization were performed using VGSTUDIOMAX, version 2.2 (Volume Graphics Inc.). Virtual sections were made in the mid-sagittal and neutral transverse planes for the vertebrae, i.e., the two reference planes for vertebral sections (see Buffrénil et al., 2008) and in the parasagittal plane for the CD13-PAL.2018.3.4 (for preservation reasons), in the coronal and sagittal planes for the humerus, and in the sagittal plane as well as in two transverse planes (proximally and at about one third of its length) for the rib.

### Phylogenetic analysis

To understand the phylogenetical affinities of the newly described genus within Mosasauroidea (*sensu* Augusta *et al*., 2022), we used the matrix of Zietlow *et al*. (2023), which is a modified version of Strong *et al*. (2020), itself adapted from the pioneer matrix of Bell (1997).

Simoes et al. (2017) and Strong et al. (2020), respectively, chose ophidiomorphs and “dolichosaurids + aigialosaurids” as outgroups for mosasauroids. In this study, however, we follow Polcyn *et al*. (2022) and Zietlow *et al*. (2023) to include the following as outgroup taxa: four species of Varanoidea, one species of Shinisauridae, and three ophidiomorph species (Appendix 1). *Ovoo gurvel* was excluded due to its incompleteness.

Our ingroup consists of 55 mosasauroid species, inclusive of the new genus described herein, the Villeveyrac mosasauroid—which is the second-most complete taxon coming from almost coeval freshwater deposits of Southern France (Garcia *et al*., 2015)—as well as six basal mosasauroids (“aigialosaurids”), three halisaurines, 23 mosasaurines, 11 plioplatecarpines, five tylosaurines, three yaguarasaurines and two tethysaurines. All taxa were scored based on their descriptions and on coding provided by Mekarski (2017) (Appendix 1).

The parsimony analysis was performed using PAUP version 4.0a169 (Swofford, 2003). According to Gauthier *et al*. (2012), multistate characters involving quantitative variation are ordered. Therefore, we designated characters 8, 19, 20, 23, 27, 29, 32, 36, 37, 41, 50, 52, 60, 75, 76, 79, 82, 83, 85, 86, 87, 89, 92, 93, 94, 100, 111, 118, 125 and 127 as ordered in our analysis. Five characters were added and one was modified (see Appendix 1). All characters were unweighted. We modified the scores for 15 taxa, especially *Pannoniasaurus* and *Tethysaurus*, based on literature and new personal observations.

The new data matrix (Appendix 2) was analysed using heuristic search algorithms. The apomorphy list was interpreted using the accelerated transformation (ACCTRAN) setting. Because three ophidiomorph taxa were included as outgroup taxa, we rooted the analysis using a paraphyletic outgroup series to provide broader sampling of putative plesiomorphic character states. Bootstrap support values were calculated and are statistically correlated with Bremer indices (e.g., bootstrap <50% corresponds to Bremer indices of 0–2). We also computed jackknife values, which are particularly informative for morphological datasets (Darlu et al., 2019). To assess data completeness and its potential impact on the resolution and robustness of the phylogenetic analyses, the percentage of missing data was calculated for each taxon (Appendix 1).

The phylogeny obtained was time-scaled (Figure 10) using the R package Strap (Bell & Lloyd, 2015) in R version 4.3.0 (2023). The first appearance datum (FAD) and last appearance datum (LAD) for each taxon were taken from the Paleobiology Database (http://paleobiodb.org/), with the exception of the new Santonian mosasauroid described herein, which was constrained (FAD = 84 Ma, LAD = 83.6 Ma) by magnetostratigraphical analyses (see Westphal & Durand, 1990), and the early Campanian Villeveyrac mosasauroid, constrained (FAD = 83.6 Ma, LAD = 80 Ma) on biostratigraphical evidence (see Garcia et al., 2015).

**Abbreviations**: CD13 - Conseil départemental des Bouches-du-Rhône, Marseille, France; MHNA - Muséum d’Histoire Naturelle d’Aix-en-Provence, Aix-en-Provence, France; MNHN - Muséum National d’Histoire Naturelle, Paris, France.

## Systematic Paleontology

Reptilia Laurenti, 1768

Squamata Oppel, 1811

Mosasauridae Gervais, 1852

Tethysaurinae Makádi *et al*., 2012

### Garamaudo bauciensis gen. et sp. nov

#### Etymology

From “*Gara au maudo*”, a Provençal expression meaning “beware the cursed one”, referring to a malevolent serpentiform creature, the Garamaudo, said to haunt local freshwaters, and first mentioned by Frédéric Mistral in *Mirèio* (1859). The specific epithet *bauciensis* means “from Bouc-Bel-Air”, the type locality.

#### Holotype

CD13-PAL.2018.3.1.1 to .3, CD13-PAL.2018.3.1.5 to .15 and CD13-PAL.2018.3.1.18 (Figures 3, 4, 5, 7), an incomplete skeleton including the incomplete posterior part of the left mandible (CD13-PAL.2018.3.1.10), seven vertebrae including five cervicals (CD13-PAL.2018.3.1.1, .3, .5, .7, .8) and two indeterminate ones (CD13-PAL.2018.3.1.2, .6), six ribs (CD13-PAL.2018.3.1.9, .12, .13, .14, .15, .18), and a right incomplete pelvis (CD13-PAL.2018.3.11), all of comparable size and found in close association.

**Figure 3.**
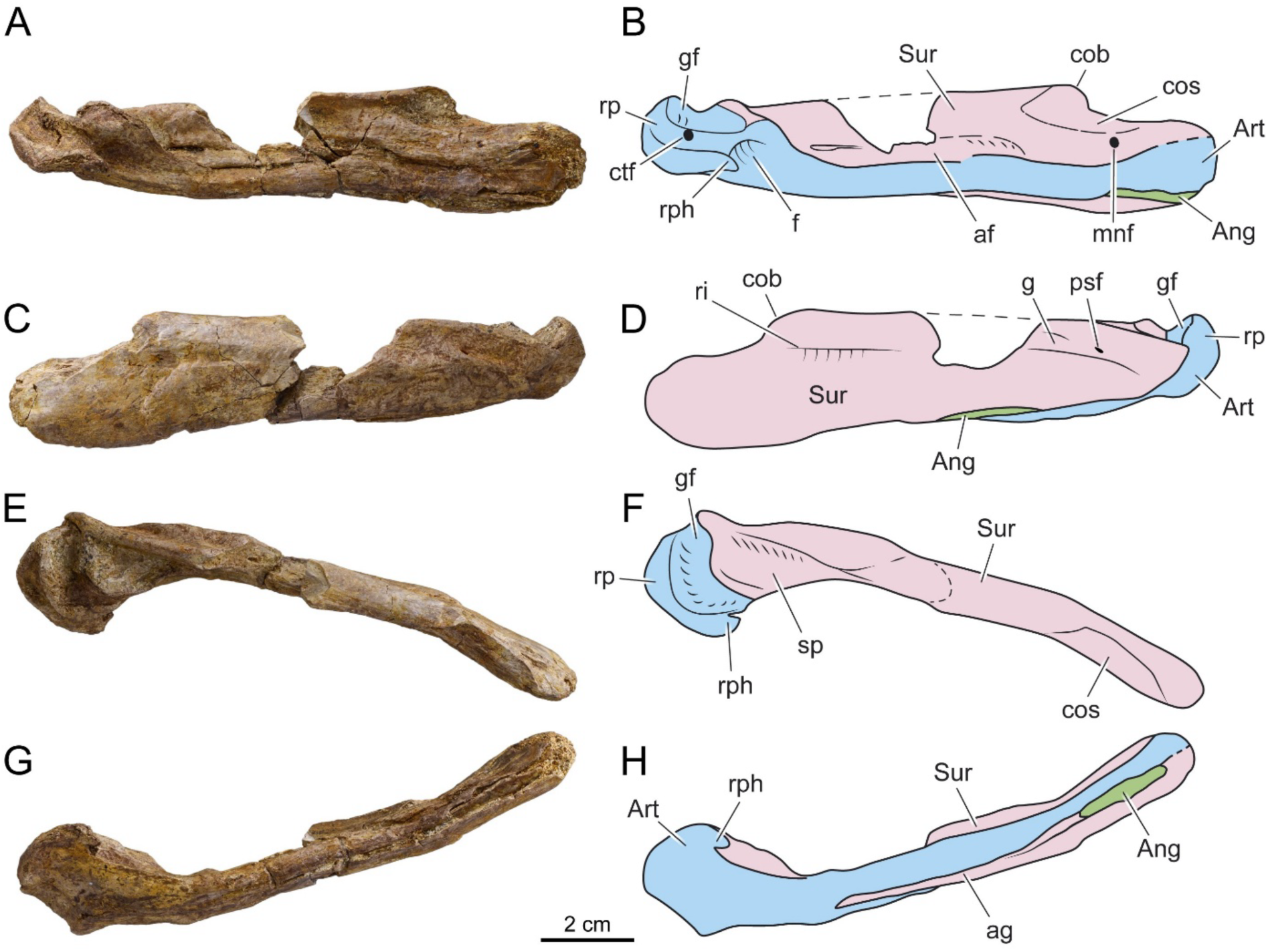
*Garamaudo bauciensis* gen. et sp. nov., mandible. Holotype (CD13-PAL.2018.3.1.10), Bouc-Bel-Air - Sousquière (Bouches-du-Rhône Department, Provence, southeastern France), Santonian. **A-H**, photographies and interpretative drawings in lateral (**A-B**), medial (**C-D**), dorsal (**E-F**) and ventral (**G-H**) views. **Abbreviations**: af, *adductor fossa*; Ang, Angular; ag, angular groove; Art, Prearticular-articular unit; cob, coronoid buttress; cos, coronoid suture; ctf, corda tympani foramen; f, fossa; g, groove; gf, glenoid fossa; mnf, mandibular nerve foramen; psf, posterior surangular foramen; ri, ridge; rp, retroarticular process; rph, retroarticular process hook; Sur, surangular; sp, surangular horizontal plateau.

**Figure 4.**
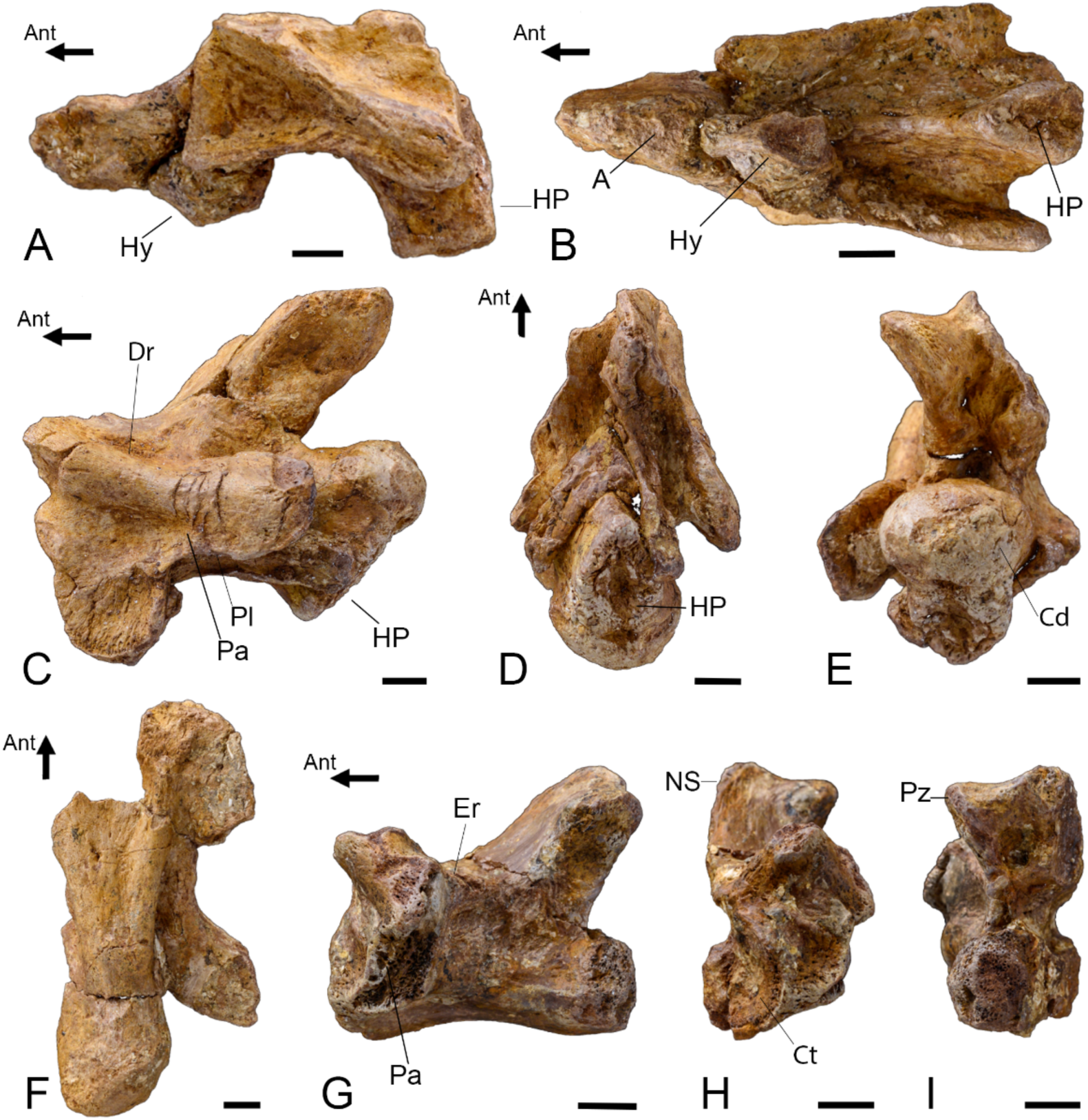
*Garamaudo bauciensis* gen. et sp. nov., vertebrae. Holotype (CD13-PAL.2018.3.1.1, .3) and referred specimens (CD13-PAL.2018.3.1.4, CD13-PAL.2018.3.4), Bouc-Bel-Air - Sousquière (Bouches-du-Rhône Department, Provence, southeastern France), Santonian. **A-B**, CD13-PAL.2018.3.1.3, Axis with part of the atlas in right lateral (**A**, mirrored) and ventral (**B**) views. **C-E**, CD13-PAL.2018.3.1.1, anterior cervical vertebra in left lateral (**C**), ventral (**D**) and posterior (**E**) views. **F**, CD13-PAL.2018.3.1.4, dorsal vertebra in ventral view. **G-I**, CD13-PAL.2018.3.4, dorsal vertebra in left lateral (**G**), anterior (**H**) and posterior (**I**) views. **Abbreviations**: A, atlas; Ant, anterior direction; Cd, condyle; Ct, cotyle; Dr, diapophyseal ridge; Er, epidiapophyseal ridge; HP, hypapophysis; Hy, hypocentrum; Ns, neural spine; NS: neural spine; Pa, paradiapophysis; Pl, parapophyseal lamina; Pz, postzygapophysis. Scale bars = 5 mm.

**Figure 5.**
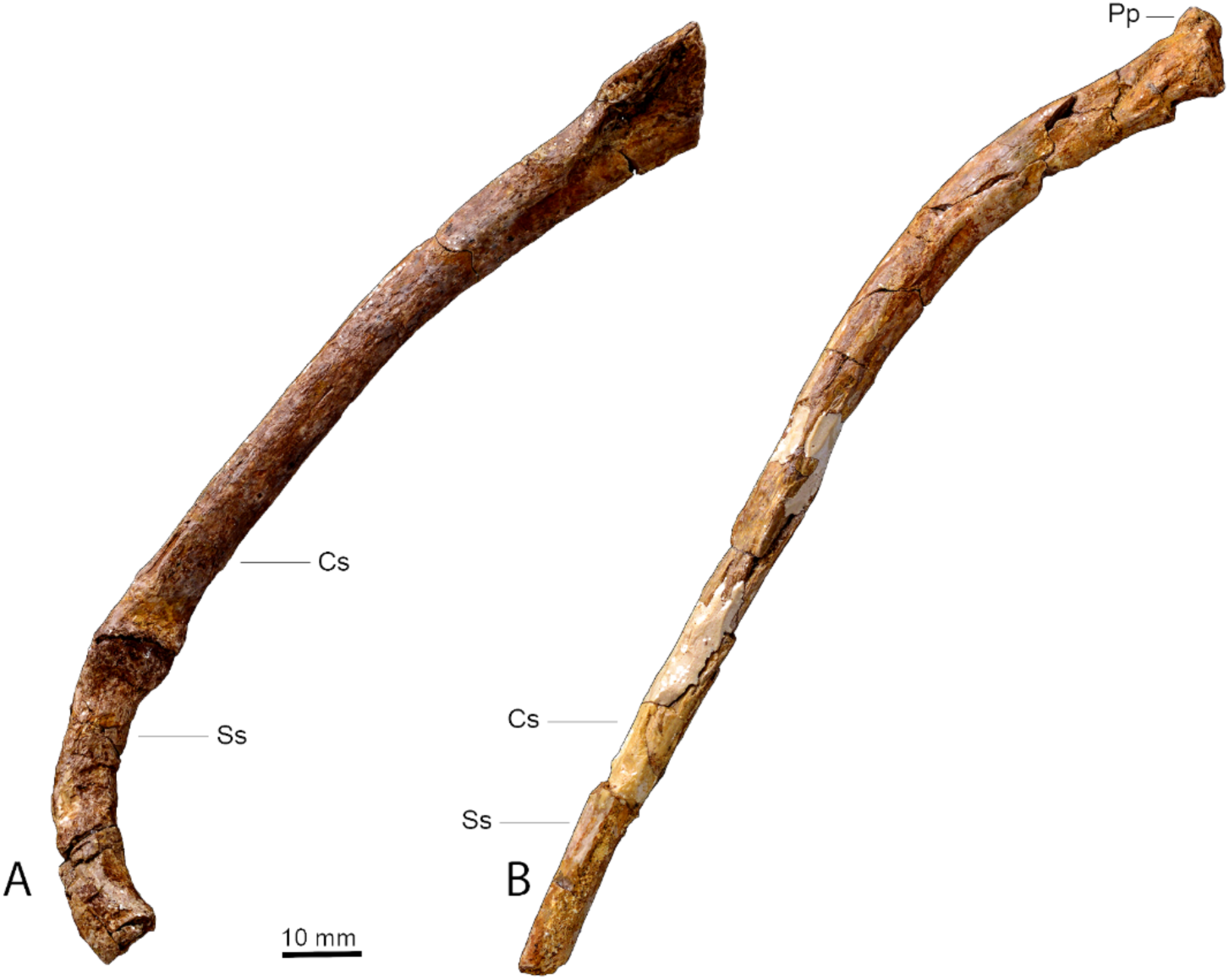
*Garamaudo bauciensis* gen. et sp. nov., ribs. Holotype (CD13-PAL.2018.3.1.9, .13), Bouc-Bel-Air - Sousquière (Bouches-du-Rhône Department, Provence, Southeastern France), Santonian. **A**, CD13-PAL.2018.3.1.9, dorsal rib, consisting of costal (proximal) and sternal segments; **B**, CD13-PAL.2018.3.1.13, posterior rib. **Abbreviations**: Cs, costal segment; Pp, posterior peduncle; Ss, sacral segment.

#### Paratype

CD13-PAL.2018.3.1.16, a right humerus, found together with the holotype but proportionally smaller (Figure 6).

**Figure 6.**
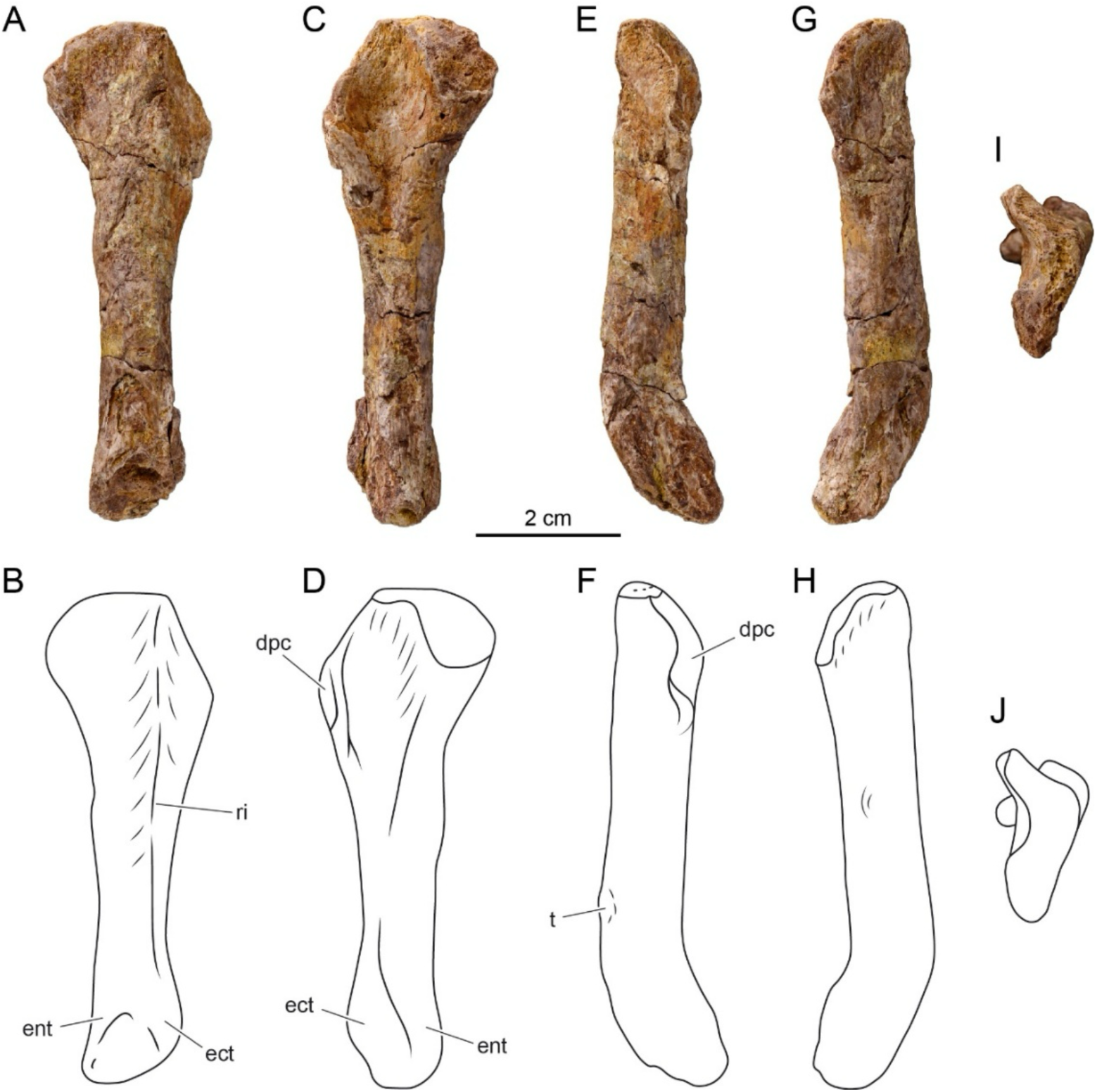
*Garamaudo bauciensis* gen. et sp. nov., right humerus. Holotype (CD13-PAL.2018.3.1.16), Bouc-Bel-Air - Sousquière (Bouches-du-Rhône Department, Provence, Southeastern France), Santonian. **A-J**, photographies and interpretative drawings in dorsal (**A-B**), ventral (**C-D**), anterior (**E-F**), posterior (**G-H**) and proximal (**I-J**) views. **Abbreviations**: ect, ectepicondyle; ent, entepicondyle; dpc, deltopectoral crest; ri, ridge; t, tubercle.

#### Referred specimens

CD13-PAL.2018.3.1.4, a dorsal vertebra, found together with the holotype but proportionally larger (centrum length estimated to be about 4.7 cm; Figure 4F); CD13-PAL.2018.3.4 (Figure 7G-I, 8E), a dorsal vertebra comparable in size with the holotype (centrum length about 2.5 cm) but found later in the same outcrop and level, about 20 m from the *locus typicus*.

**Figure 7.**
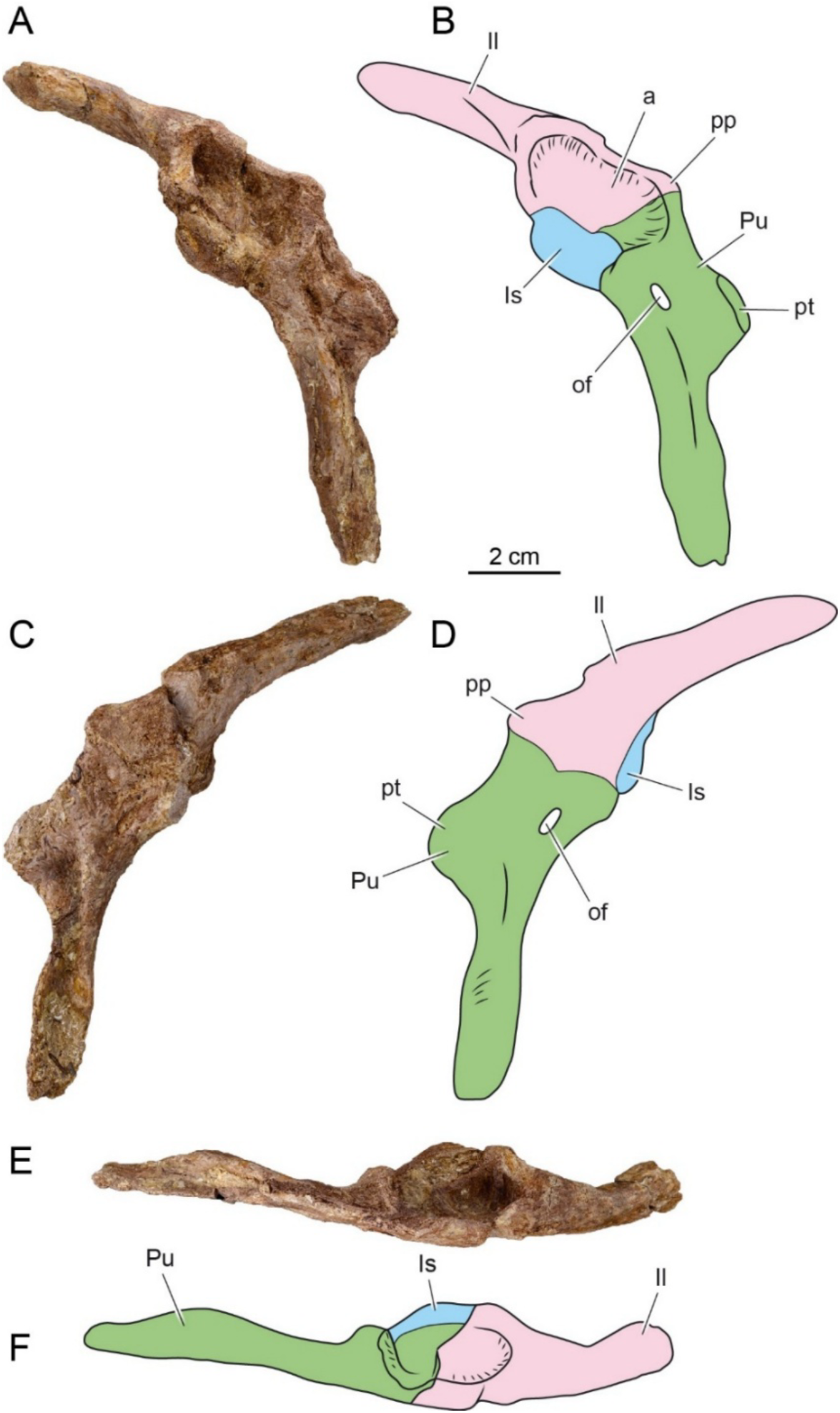
*Garamaudo bauciensis* gen. et sp. nov., pelvic girdle. Holotype (CD13-PAL.2018.3.1.11), Bouc-Bel-Air - Sousquière (Bouches-du-Rhône Department, Provence, Southeastern France), Santonian. **A-F**, photographies and interpretative drawings in lateral (**A-B**), medial (**C-D**) and dorsal (**E-F**) views. **Abbreviations**: a, acetabulum; Il, Ilium; Is, Ischium; of, obturator foramen; pp, preacetabular process; Pu, Pubis; pt, pubic tubercle.

#### Geographical and stratigraphical occurrences

Bouc-Bel-Air - Sousquière outcrop (BBAS), near Aix-en-Provence, Bouches-du-Rhône Department, Provence, southeastern France; Level 99, Valdonnian local facies, Santonian (Durand & Guieu, 1980).

#### Diagnosis

*Garamaudo bauciencis* is a tethysaurine mosasaurid characterized by the unique combination of the following characters (autapomorphies indicated by *) : posterior mandibular unit (PMU) long, slender, and rectangular, with glenoid fossa made mainly by the articular (shared with basal mosasauroids), and strongly curved inward in dorsal/ventral view*; surangular with a medial horizontal triangular plateau located just anteriorly to the anterior border of the glenoid fossa*; retroarticular process oriented obliquely about 45° to the long axis of the PMU, extremely short (half the length of the glenoid fossa), with a convex semi-lunate posterior margin extending antero-medially and ending as an anteriorly oriented hook*; adductor fossa of the surangular open along its length, being located dorsally and parallel to the surangular/prearticular-articular suture*; dorsal ribs laterally compressed, very long and straight, being only curved at the proximal third of their length, indicating a dorsoventrally high and laterally compressed ribcage (shared with *Mesoleptos*); humerus long, slender, and straight, with epiphyses oriented almost perpendicular to one another (shared with *Varanus* but shorter), and possibly reduced* (see Discussion); pelvic girdle with ilium, ischium, and pubis firmly fused to form an oval acetabulum (shared with basal mosasauroids); ilium posteriorly oriented (shared with basal mosasauroids); pubis with a very long and slender anterior process, in alignment with the body longitudinal axis*.

## Comparative description

### General preservation

The vertebrae are not well preserved (none is complete) and exhibit various lateral or dorsal compressions, breakages, and frequent abrasion, perforation and/or selachian feeding marks (NB pers. obs.) on their external surfaces. Conversely, the other bones (posterior mandibular unit, humerus, pelvis) remain relatively complete and undistorted, with their external surfaces relatively well preserved, limiting the risks that some observed characteristics are linked to taphonomical biases. However, many of the sutures are poorly visible, especially on the PMU and on the pelvis, which is why interpretative drawings are provided.

The state of preservation of the bones indicates that they may have been transported a short distance prior to their burial (explaining why some bones remain relatively complete and not abraded or compressed). The disarticulated and broken state of some of the bones suggests that the transportation may have resulted from a high-energy flooding episode. This interpretation agrees with the proposed paleoenvironment, which is a freshwater lake periodically influenced by flooding and shallowing events (see Discussion). In addition, the occurrence of possibly four individuals belonging to the same taxon suggests that this basal mosasaurid was probably autochthonous or parautochthonous to this freshwater environment, as already observed in several Upper Cretaceous continental outcrops from Europe (see Introduction and Discussion).

The poorly visible sutures on both the PMU and pelvic girdle, and the absence of spongy parts on the outer surface of the bones as well as the relatively high tightness (high number of thin trabeculae) of the spongiosa of the vertebrae suggest a mature ontogenetic stage (Houssaye & Bardet, 2013).

### Cranial skeleton

Only the left PMU (CD13-PAL.2018.3.1.10; abbreviated as PMU from now on) is preserved, including the almost complete surangular and prearticular-articular, and a very small part of the angular (Figure 3). The coronoid, which is usually loosely “saddled” above the surangular and prearticular, has been lost postmortem. The PMU has the shape of a long and low rectangle in lateral/medial views (length about 12.8 cm, median height about 2.3 cm) (Figure 3A-D) and is curved inward in dorsal/ventral view (Figure 3E-H). The ventral margin of the PMU is slightly sigmoid in lateral/medial view, convex anteriorly, concave in its median part, and convex again posteriorly (Figure 3A-D).

The surangular is an elongated and low rectangular bone (length about 11.8 cm) that occupies most of the lateral surface of the PMU (Figure 3A-B), except its posterior part occupied by the articular. Its anterior portion, though not well preserved, is probably not vertical but slightly obliquely oriented in lateral view. The lateral surface of the bone is gently convex dorsoventrally and bears a marked longitudinal straight ridge (for attachment of the *pseudotemporalis superficialis* and *adductor mandibulae* muscles), located slightly above the mid-height of the bone and rising slightly forward (“ri”, Figure 3D). Anteriorly, this ridge is bordered ventrally by a shallow groove; posteriorly the ridge is rugose and bordered by an oval concavity and an oval foramen (“psf”, Figure 3D), nestled in a shallow groove that terminates just ventral to the anterior margin of the glenoid fossa. Ventral to the posterior part of the ridge, the surangular is very rugose. The ventral suture of the surangular with the prearticular-articular is straight and located on the ventral margin of the mandible so that the prearticular-articular is poorly exposed in lateral view (Figure 3A-B). Posteriorly, this suture curves abruptly dorsally at an angle of about 50° to the posterolateral corner of the glenoid fossa, where the surangular forms a blunt, laterally projecting corner. From this corner, the surangular gently rises in a straight line anteriorly for most of the length of its dorsal margin, before curving down abruptly to form a concave articulation with the coronoid. In dorsal view, from the antero-lateral sharp corner of the glenoid fossa, the surangular curves first medially and then anteriorly, forming a sigmoid elevated dorsal lip that forms the anterior border of the glenoid fossa and its only participation to it (Figure 3E-F). This lip continues anteriorly on the medial surface of the bone and forms a unique concave triangular plateau roughly horizontally oriented (“sp”, Figure 3F), before the bone becomes laterally compressed along its length. In medial view, the suture with the prearticular-articular is concave from the anterior border of the glenoid fossa, then runs horizontally along most of its length before sloping dorsally in its anterior part (Figure 3C-D). This suture is poorly visible and is located on the lower third of the height of the PMU, meaning that the participation of the surangular in the medial surface of the PMU is much larger than that of the prearticular-articular. The shallow *adductor fossa* (“af”, Figure 3B) is located dorsally and parallel to the surangular/prearticular-articular suture. It is unique in being open (i.e., not covered anteriorly by a thin descending blade of bone), and, as such, it extends for most of the length of the surangular. The region where the mandibular foramen should have occurred is broken. Anterodorsally, the suture for the coronoid is long (“cos”, Figure 3B,F), occupying about a third the length of the PMU, and shallow, showing that the medial coronoid wing did not extend far ventrally. The coronoid buttress is also poorly developed (“cob”, Figure 3B,D).

The angular is possibly represented by two tiny thin blades of bone located between the surangular and the prearticular-articular: one anteriorly, about 2.5 cm long, visible in medial view; the other is visible laterally around the middle ventral surface of the PMU (Figure 3A-D). The ventral surface of the PMU bears a shallow, slender longitudinal groove for the articulation of the angular that extends very far posteriorly (for about three-quarters of its total PMU length) where it tapers to a sharp, thin extremity (“ag”, Figure 3H). Preserved in its entirety, the angular would have given the ventral margin of the PMU a straighter aspect.

The prearticular and articular are fused into a single bone without any visible suture, as in all squamates (Russell, 1967; Zietlov *et al*., 2023). The unit occupies the entire length of the PMU (that is 12.8 cm) and half of its medial surface (Figure 3C-D), but remains obscured in lateral view (Figure 3B). In medial view, the prearticular-articular is a horizontal slender bone that expands posteriorly at the level of the glenoid fossa and retroarticular process. The glenoid articulation retains the plesiomorphic condition, whereby it is formed mainly by the articular, with the surangular participating only in its anterior margin. The glenoid is slightly concave and transversally oriented, about twice as wide as long, and reniform, with a sigmoid anterior margin and a concave posterior one (Figure 3C-F). Both anterior and posterior margins are elevated rugose “lips”, whereas the lateral and medial margins are flat. The fan-shaped retroarticular process is oriented obliquely at about 45° to the long axis of the PMU (Figure 3E-H). It is unique both in being extremely short (half the length of the glenoid fossa) and in bearing an evenly convex semi-lunate posterior margin. This posterior margin extends antero-medially to form an anteriorly oriented hook, creating a medial fossa ventral to the glenoid fossa (“rph”, Figure 3B,F,H). The retroarticular process bears a large, oval foramen for the corda tympani (“ctf”, Figure 3B), located just ventral to the posteromedial corner of the glenoid fossa.

The PMU of *Garamaudo bauciensis* is typical of squamates but differs greatly from that of snakes (whose surangular and articular/prearticular are fused into a single complex; e.g., Lee 1997) and from ophidiomorphs (whose articular is usually visible in lateral view between the angular and the surangular; e.g., Pierce & Caldwell, 2004). It also differs from that of derived mosasaurids, in which the PMU is proportionally stouter, shorter, and more diamond shaped in lateral view (Russell, 1967). The PMU of *G. bauciensis* is comparable to that of basal mosasauroids (e.g., *Aigialosaurus, Haasiasaurus*, tethysaurines and yaguarasaurines) in having a long and slender rectangular shape, and in possessing a glenoid articulation composed mainly by the articular, with a reduced contribution of the surangular (DeBraga & Carroll, 1993).

### Axial skeleton

#### Vertebrae

All vertebrae are procoelous and not pachyostotic (Figure 4). Based on the broken parts available, it appears that prezygapophyses project strongly anteriorly, extending well beyond the anterior border of the cotyle (Figure 4C, G).

#### Cervical vertebrae

CD13-PAL.2018.3.1.3 consists of an articulated partial atlas-axis (Figure 4A-B). The axis has a long hypapophysis that projects strongly ventrally, terminating in an anteroposteriorly directed ovoid facet that is almost horizontal. Anteriorly, the axis intercentrum articulates ventrally with both the anteriormost part of the centrum, through an anteroventral facet, and the atlas, through a postero-ventral facet. It seems to end in a small, anteroposteriorly elongated ovoid facet that points ventrally and slightly posteriorly. Paradiapophyses are very short dorsoventrally and very posteriorly located, with marked dorsal and ventral ridges. The neural canal is much thinner than the condyle.

CD13-PAL.2018.3.1.1 is an anterior cervical vertebra (Figure 4C-E), as shown by the well-developed hypapophysis, continuous with the centrum that ends below the condyle in an ovoid concave facet elongated dorsoventrally and oriented strongly posteriorly and ventrally (Figure 4C-D). The hypocentrum is not fused. The centrum is elongated. The condyle is ovoid (Figure 4E), wider than tall (dorsoventrally compressed), and there is no precondylar constriction. The cotyle-condyle axis is horizontal. The paradiapophyseal facet is dorsoventrally short and projects strongly posteriorly. A marked anterodorsal diapophyseal ridge links its dorsal border to the prezygapophysis. Anteriorly, the ventral parapophyseal lamina (extending anteroventrally from the paradiapophyses; see Houssaye *et al*., 2011) extends well beyond the ventral border of the cotyle. The preserved postzygapophysis has a large facet directed lateroventrally (Figure 4E). The posterior aspect suggests the occurrence of a zygantrum, although no zygantral facet is visible, apparently with zygosphenal foramina. The neural canal appears cylindrical, with a flattened ventral border and a trilobate aspect formed by the two ‘‘ridges’’ running along the lateral walls. The canal is clearly narrower and shorter than the condyle (Figure 4E). The short, flat-topped neural spine rises from about a third of the length of the centrum and does not extend beyond the condyle (Figure 4C).

CD13-PAL.2018.3.1.8 is a broken anterior cervical vertebra, as shown by the probably ventral extension of the paradiapophyses and the hypapophysis with a facet facing most likely posteriorly.

CD13-PAL.2018.3.1.6 and CD13-PAL.2018.3.1.7 are parts of the same broken vertebra. These represent a more posterior cervical vertebra with a taller cotyle, probably more circular. The paradiapophyses extend slightly below the centrum and become very tall posteriorly. The posterior part of the centrum is not preserved, preventing us from determining the possible occurrence of a hypapophysis.

CD13-PAL.2018.3.1.5 consists of very partial, articulated cervical vertebrae with paradiapophyses projecting far beyond the ventral border of the centrum, and a prezygapophysis projecting far anteriorly.

#### Dorsal vertebrae

CD13-PAL.2018.3.4 is an anterior dorsal vertebra (Figure 4G-I). The paradiapophyses do not extend below the ventral border of the centrum (Figure 4H) and are very anteriorly located. They are subvertical and extend up to the base of the prezygapophyses (Figure 4G). The epidiapophyseal ridge is well marked. In dorsal view, the narrowest part of the interzygapophyseal constriction occurs posteriorly. The centrum is relatively shorter than in CD13-PAL.2018.3.1.4. and more triangular in ventral view. The posterior border of the neural arch is concave, and the arch is thick (Figure 4I), while the neural spine appears rather short. It extends from below the paradiapophyses, is shorter than in CD13-PAL.2018.3.1.1., and does not extend beyond the condyle (Figure 4G). The straight or oblique nature of the cotyle-condyle axis cannot be determined. One zygantral foramen is visible.

CD13-PAL.2018.3.1.4 is a large elongated vertebra with an ovoid condyle, dorsoventrally compressed and a long and flat centrum is reminiscent of posterior dorsals (Figure 4F).

#### Indeterminate vertebra

CD13-PAL.2018.3.1.2 is extremely partial and only shows that the prezygapophyses are oriented antero-laterally, and that the cotyle is ovoid.

The vertebrae of *G. bauciensis* are procoelous, as in most other squamates (Hoffstetter & Gasc, 1968). They display features of Varanoidea *sensu* Lee (1997), i.e., Varanidae, Helodermatidae and Pythonomorpha: a hypapophysis of central origin with an articulated distal element, and the posterior position of the narrowest part of the interzygapophyseal constriction. The likely occurrence of a zygantrum suggests that a zygosphene/zygantrum articulation was probably present. It characterizes Pythonomorpha Cope, 1869 *sensu* Lee (1997), i.e., Mosasauroidea and Ophidiomorpha *sensu* Palci & Caldwell (2007; i.e., stem Ophidia [*Adriosaurus*, *Aphanizocnemus*, Dolichosauridae] and Ophidia), that is the clade including the most recent ancestor of mosasaurs, snakes and all its descendants (node-based definition from Caldwell, 2006). The general morphology of the vertebrae is clearly distinct from that of snakes, although the diagnostic characters are not observable on this limited material. It differs also from that of hydropedal mosasauroids and of the plesiopedal and hydropelvic mosasauroid *Dallasaurus* whose centrum is notably more cylindrical and less flat (e.g., Russell, 1967; Bell & Polcyn, 2005). The vertebrae appear very similar to those of freshwater plesiopedal tethysaurines, which are the Villeveyrac taxon from southern France (Garcia et al., 2015) and *Pannoniasaurus* from Hungary (Makádi *et al*., 2012).

#### Ribs

Six ribs are preserved, of which two are almost complete. They are very elongated and straight, curving only at the proximal third of their length (Figure 5). As a consequence, the ribcage in cross section would have been deep and laterally compressed. The ovoid, medio-laterally compressed head bears a small posterior peduncle.

CD13-PAL.2018.3.1.9 is a dorsal rib consisting of the proximal costal segment and the distal sternal segment, the latter being calcified cartilage in life. The sternal segment is curved medially.

CD1 3-PAL.2018.3.1.13 is a more posterior rib. It is longer than CD13-PAL.2018.3.1.9, measuring about 15.0 cm in a straight line and 15.2 cm along its curvature (from the most proximal to the most distal points preserved). The length is less than 11.0 cm (11.5 cm along its curvature) if only the dorsal segment is considered.

Other proximal portions of ribs show a morphology consistent with the above.

The rib morphology is consistent with that of the stem-Ophidia *Mesoleptos* and of plesiopedal and plesiopelvic mosasauroids (e.g., *Aigialosaurus bucchichi*; *Komensaurus*; Caldwell & Palci, 2007; Dutchak & Caldwell, 2009).

### Appendicular skeleton

The appendicular skeleton is represented by a right humerus (paratype) and the right half of the pelvis (holotype).

#### Anterior limb

The humerus (CD13-PAL.2018.3.1.16) is long (preserved length about 7.0 cm), straight, and slender. The long diaphysis contributes to a third of the total length (Figure 6A-H), and the proximal and distal extremities are nearly perpendicular to one another (Figure 6I-J). The diaphysis is slightly flattened dorsoventrally, except its median part, which is thick (about 1.0 cm wide) and roughly rounded in cross-section. The proximal articular surface is oval, about 1.9 cm wide, rugose, and slightly posteriorly deflected from the longitudinal axis of the diaphysis. The deltopectoral crest is a thin but protrusive blade, developed along the proximal third of the humerus (“dpc”, Figure 6C-F). It is not separated from the proximal articular head by a deep groove, like it is in *Varanus*, and forms a 115° angle with the head, creating a shallow concave triangular area between these structures (Figure 6C-D). The dorsal surface of the humerus is strongly convex and bears a sharp longitudinal ridge that runs from the posterior corner of the proximal articulation surface to the posterior ectepicondyle zone (“ri”, Figure 6A-B). It probably corresponds to the insertion area of the *M. latissimus dorsi* (cf. Russell, 1967). The distal epiphysis is broken, but the fragmentary entepicondyle and ectepicondyle (“ect” and “ent”, Figure 6A-D) show that it was probably expanded. A small tubercle is preserved on the dorsal surface of the humerus along its distal third (“t”, Figure 6E-F), as seen in *Varanus* and basal mosasauroids (DeBraga & Carroll, 1993). While long and slender, the humerus is considerably shorter than the PMU; in extant varanids and plesiopedal mosasaurids, it is about the same length (NB, pers. obs).

The humerus *de facto* excludes this taxon from snakes. It is clearly plesiopedal in nature, being long and slender, with a long diaphysis making about 1/3 of the total length, and proximal and distal extremities oriented almost perpendicular to each other, like in *Varanus*. However, the bone is flatter, shorter and with a stouter, rounded epiphysis than in *Varanus*, being reminiscent - though longer and slender - of that of plesiopedal mosasauroids, such as *Aigialosaurus* and *Carsosaurus* (DeBraga & Carroll, 1993) or the mosasaurine *Dallasaurus* (Bell & Polcyn, 2005). It differs greatly from the humerus of hydropedal mosasauroids, which is generally very reduced in length compared to its width (Russell, 1967). To sum up, the humerus is intermediate in shape and proportions between varanids and basal plesiopedal mosasauroids.

#### Pelvic girdle

The pelvis (holotype, CD13-PAL.2018.3.11) is about 14.0 cm long. It includes the almost complete ilium and pubis and only the acetabular portion of the ischium (Figure 7). The three bones are firmly fused together at the acetabulum, which is deeply concave and roughly oval. The suture between the three bones is clearly visible in both lateral and medial views (Figure 7A-D).

The ilium is about 8.0 cm long but its posterior extremity is incomplete, so that it could have been about the same length as the pubis. This incompleteness precludes determination of whether a distal articulation for the sacrum was present or not. The ilium is a robust rod of bone that is posterodorsally oriented. Its contribution to the acetabulum is the largest of the three pelvic bones, forming the entire dorsal portion (“a”, Figure 7A-B). The ventral articulations of the ilium for both the pubis anteriorly and the ischium posteriorly are obliquely oriented to each other at about 120°, forming an open V shape (Figure 7A-B). The anterior preacetabular process is a small rounded tongue of bone, laterally compressed and anteriorly oriented, slightly overlapping the pubic articulation (“pp”, Figure 7A-D). Posteroventrally, the ilium abuts the ischium along a reinforced, rounded portion of the latter.

The pubis is about 8.3 cm long and almost flat. It contributes to the anteroventral part of the acetabulum. Here, it forms an oblique straight suture with the ilium and a shorter, posteriorly concave suture with the ischium (“a”, Figure 7A-D). The proximal part of the pubis is expanded, with a large rectangular pubic tubercle, which is mostly dorsally and only slightly laterally oriented (“pt”, Figure 7A-D). A large and oval obturator foramen occurs roughly in the medio-ventral part of the bone (“of”, Figure 7A-D). The anterior process of the pubis is unique in being very long and slender, triangular in section, and oriented in the same plane as the longitudinal axis of the body (Figure 7E-F). In lateral view this anterior process is curved from the horizontal, forming a bowed structure ending in a vertical tip. Also, in lateral view, its dorsal surface is sigmoid, being strongly concave anterior to the pubic tubercle, then convex, finally straight, whereas its ventral surface is regularly concave along its length.

The ischium is represented only by the acetabular contribution, having a long and oblique suture with the ilium and a shorter, posteriorly concave one with the pubis (Figure 7A-D). It is not possible to determine if, like the pubis, the ischium occurred in the same plane as the ilium and pubis or was medially oriented to meet its left counterpart in a median suture (e.g., DeBraga & Carroll, 1993).

The pelvis also excludes this taxon from snakes. The pelvis of the few snakes with a hindlimb, when preserved, is a relatively much smaller triradiate structure, but poorly known and described, which limits comparisons (Rieppel *et al*., 2003; Houssaye *et al*., 2011; Palci *et al*., 2013). In possessing three firmly fused bones forming a large oval acetabulum, as well as a posteriorly oriented ilium, the pelvis looks very much like that of varanids and basal plesiopelvic mosasauroids (“aigialosaurids”, tethysaurines, situation unknown in yaguarasaurines) (DeBraga & Carroll, 1993) and strongly differs from hydropelvic mosasaurids, where the pelvic bones are unfused, and the ilium is anteriorly oriented (Russell, 1967). On the contrary, like hydropelvic mosasaurids, the ilium bears a small and rounded preacetabular process abutting on the pubis, whereas the pubis has a pubic tubercle mostly dorsally oriented, and a long and slender anterior process parallel to the body long axis. In varanids, the preacetabular process of the ilium is longer and distinct from the main shaft of the bone, the tubercle of the pubis is rounded and mostly ventrolaterally oriented, and its anterior process, though also long, is oriented perpendicular to the body long axis to meet its counterpart in a median suture (DeBraga & Carroll, 1993). In basal mosasauroids, the preacetabular process of the ilium is also longer, whereas the anterior process of the pubis is shorter and more fan shaped distally, and appears oriented parallel to the long axis of the body (e.g., DeBraga & Carroll, 1993). As a whole, the pelvis is unique in exhibiting a mosaic of characters present in both terrestrial varanids, plesiopelvic, and hydropelvic mosasauroids.

### Microanatomy

The vertebrae show a relatively thick layer of compact cortex, slightly thicker in the axis (Figure 8A) than in the anterior cervical and dorsal vertebrae (Figure 8C). Inner compactness is high as compared to in modern lizards. Vertebral microanatomy is far from the typical tubular inner organization of squamates, with one outer layer of compact bone surrounding the bone and another surrounding the neural canal, and a few trabeculae connecting them (Houssaye *et al*., 2010). Even the endochondral region shows only small cavities. However, larger cavities linked to active resorption are visible in the periosteal region. The hypapophysis is less compact than the rest of the vertebra. Some simple, radially oriented vascular canals are observed in the cortex in transverse sections, notably in CD13-PAL.2018.3.4 (Figure 8B, D, E; much less visible in the longitudinal ones).

**Figure 8.**
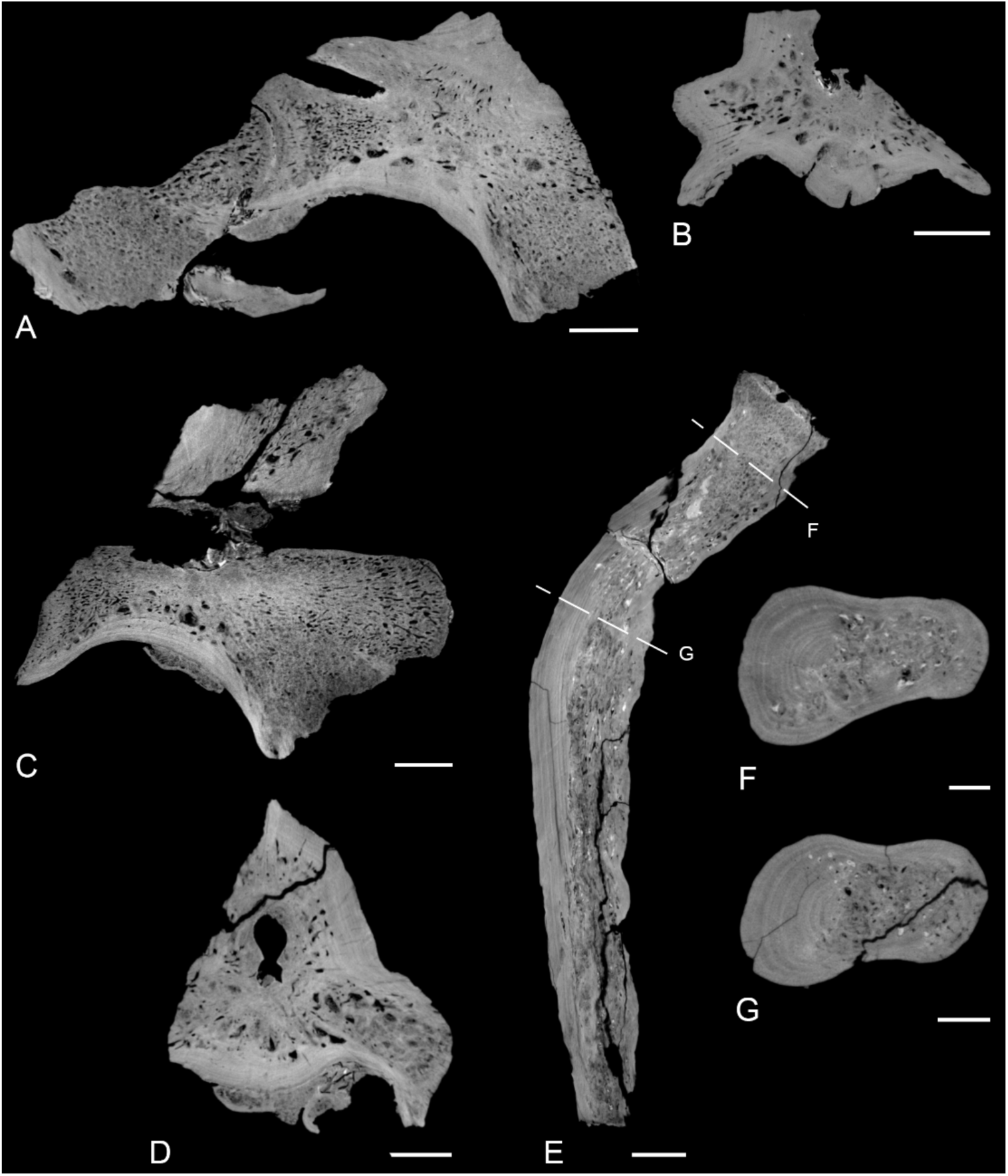
*Garamaudo bauciensis* gen. et sp. nov., vertebral and rib microanatomy. Holotype (CD13-PAL.2018.3.1.1, .3) and refered specimen (CD13-PAL.2018.3.4), Bouc-Bel-Air - Sousquière (Bouches-du-Rhône Department, Provence, Southeastern France), Santonian. **A-D**, Vertebral virtual sections of CD13-PAL.2018.3.1.3 (**A-B**) and CD13-PAL.2018.3.1.1 (**C-D**), in mid-sagittal (**A, C**) and transverse (near the growth center; **B, D**) sections. **E-G**, Rib CD13-PAL.2018.3.1.13 longitudinal (**E**) and transverse (**F-G**; see sectional planes on **E**) sections. Scale bars = 5mm (**A-E**) and 2 mm (**F-G**).

**Figure 9.**
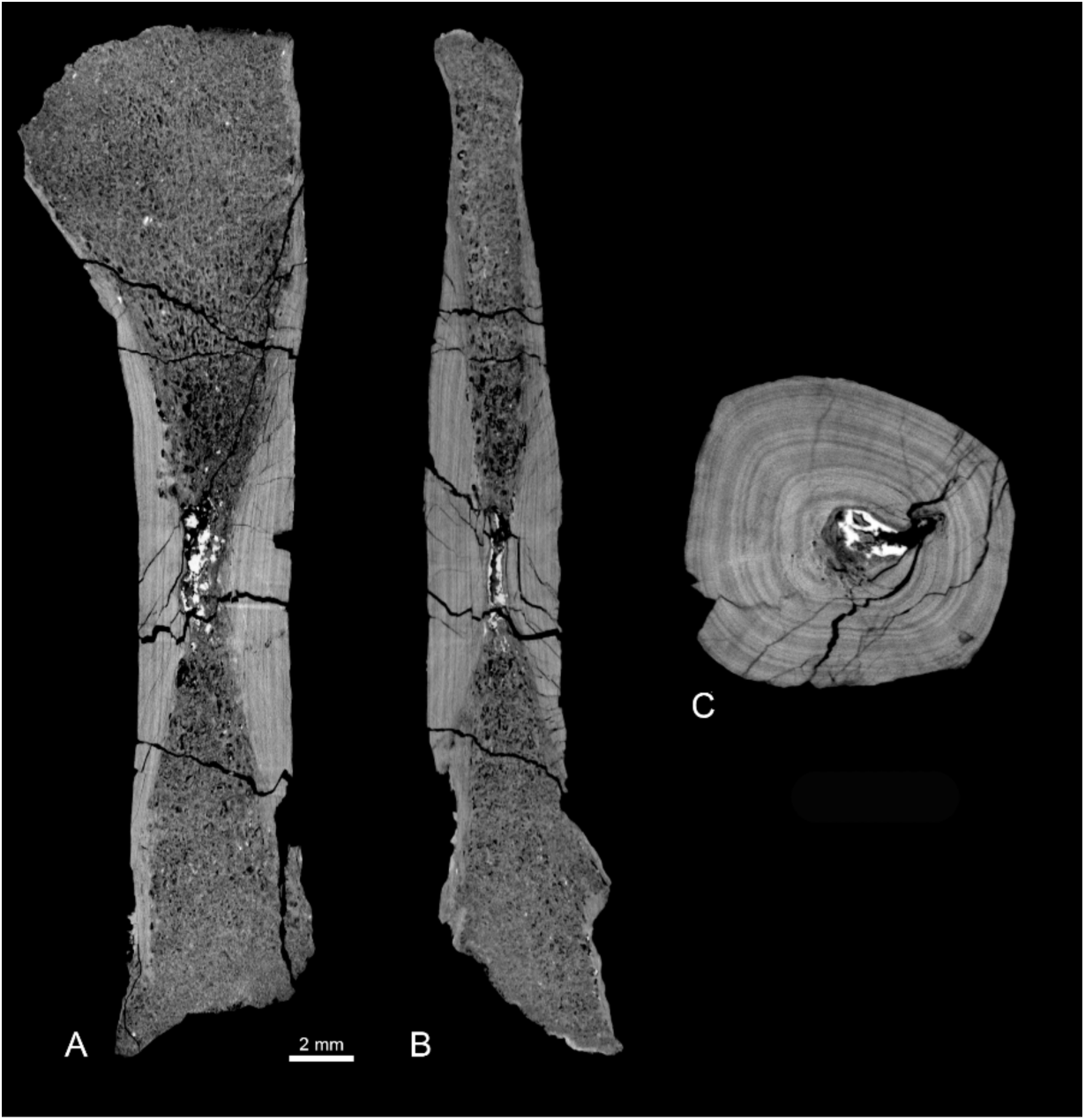
*Garamaudo bauciensis* gen. et sp. nov., humerus microanatomy. Paratype (CD13-PAL.2018.3.1.16), Bouc-Bel-Air - Sousquière (Bouches-du-Rhône Department, Provence, Southeastern France), Santonian. **A-C**, coronal (**A**), sagittal (**B**) sections, and transverse section around the growth center (**C**).

The rib shows a thick cortex surrounding a compact spongiosa and no open medullary cavity (Figure 8F-G). There is a strong asymmetry, with a much thicker cortex dorsally (Figure 8F). The thickness increases progressively from the proximal part to about a third the length of the rib, then progressively decreases.

The longitudinal sections of the humerus show that compact cortex distribution delimits a clear hourglass shape. Its maximal compactness, at the growth center, appears close to the mid-diaphysis (Figure 8A-C). There is a small medullary cavity in the core of the diaphysis, and the rest of the medullary area is filled by a spongiosa whose tightness decreases toward the core of the bone. As a consequence, the bone is very compact. Primary cortical bone clearly displays growth marks, and some radial simple vascular canals are observed.

The vertebrae, rib, and humerus of *Garamaudo bauciensis* all clearly show osteosclerosis (as compared to modern squamates; Houssaye *et al*., 2010), with an inhibition of primary bone resorption, as shown by the relatively high thickness of primary compact cortex. The primary nature of these deposits is clearly indicated by the growth marks and vascularization when visible.

The microanatomy allows this taxon to be differentiated from several pythonomorphs as follows: 1) The apparent absence of pachyostosis on the available vertebrae differentiate it from most ophidiomorphs that display at least some vertebrae with a bloated aspect (Houssaye, 2013a); 2) The occurrence of osteosclerosis in the vertebrae clearly distinguishes this taxon from the other ophidiomorphs (*Dolichosaurus* and *Coniasaurus*) that display a “classical” tubular structure (Houssaye, 2013a), from the lizard from Lo Hueco (Houssaye *et al*., 2013a) that shows compactness values within the range, although among the highest, of modern squamates and a tubular structure in transverse sections, and from hydropedal and hydropelvic mosasauroids with their tight trabecular network (Houssaye & Bardet, 2012). Compactness is, however, much lower than in *Carentonosaurus*, *Pachyvaranus*, the pythonomorph from Touraine, *Mesoleptos*, and *Haasiasaurus* (Houssaye, 2013a). It also distinguishes this taxon from the plesiopedal and plesiopelvic mosasauroids *Tethysaurus* and *Dallasaurus* that display a rather loose and heterogeneous spongious organization inside their vertebrae, with a higher density in the periosteal region, but much less dense than in *Garamaudo* (Houssaye, 2008). Similarly, the osteosclerosis in the ribs differs from the much lighter structure observed in the ribs of hydropedal and hydropelvic mosasauroids, and from the open medullary cavity observed in the ophidiomorphs *Kaganaias* and *Dallasaurus*. It also differs from the more extensive osteosclerosis in *Carentonosaurus* and the ophidiomorph *Pontosaurus.* The humerus is strongly osteosclerotic, to a similar or higher extent than in *Dallasaurus* (the only plesiopedal mosasauroid for which a humerus section is available, to our knowledge; Houssaye *et al*., 2013b).

The vascularization of the vertebrae consists of simple canals. It is limited to a few canals in the cervical vertebrae, with much more extensive vascularization occurring in the dorsal vertebra CD13-PAL.2018.3.4.

Only a few simple vascular canals occur in the largest modern terrestrial squamates, and all others do not show any vascularization (Buffrénil *et al*., 2008; Houssaye *et al*., 2010). In *Garamaudo*, vascularization is in fact similar to that observed in some pythonomorph lizards such as the marine varanoid *Pachyvaranus* (Buffrénil *et al*., 2008), the “pythonomorph from Touraine” (Houssaye *et al*., 2010), or the ophidiomorph *Carentonosaurus* (Houssaye *et al*., 2008). However, the degree of vascularization is clearly lower than in mosasaurids (Houssaye & Bardet, 2012), including the plesiopedal *Dallasaurus*. Indeed, whereas primary osteons appear absent in *Garamaudo*, they are present in mosasaurids.

## Phylogenetic Analysis

The analysis recovered 537,600 most parsimonious trees (MPTs) with 632 steps each. A strict consensus tree was generated (Consistensy Index = 0.2574, Homoplasy Index = 0.7426, Retention Index = 0.6638) and transposed as a time-scaled tree (Figure 10).

**Figure 10.**
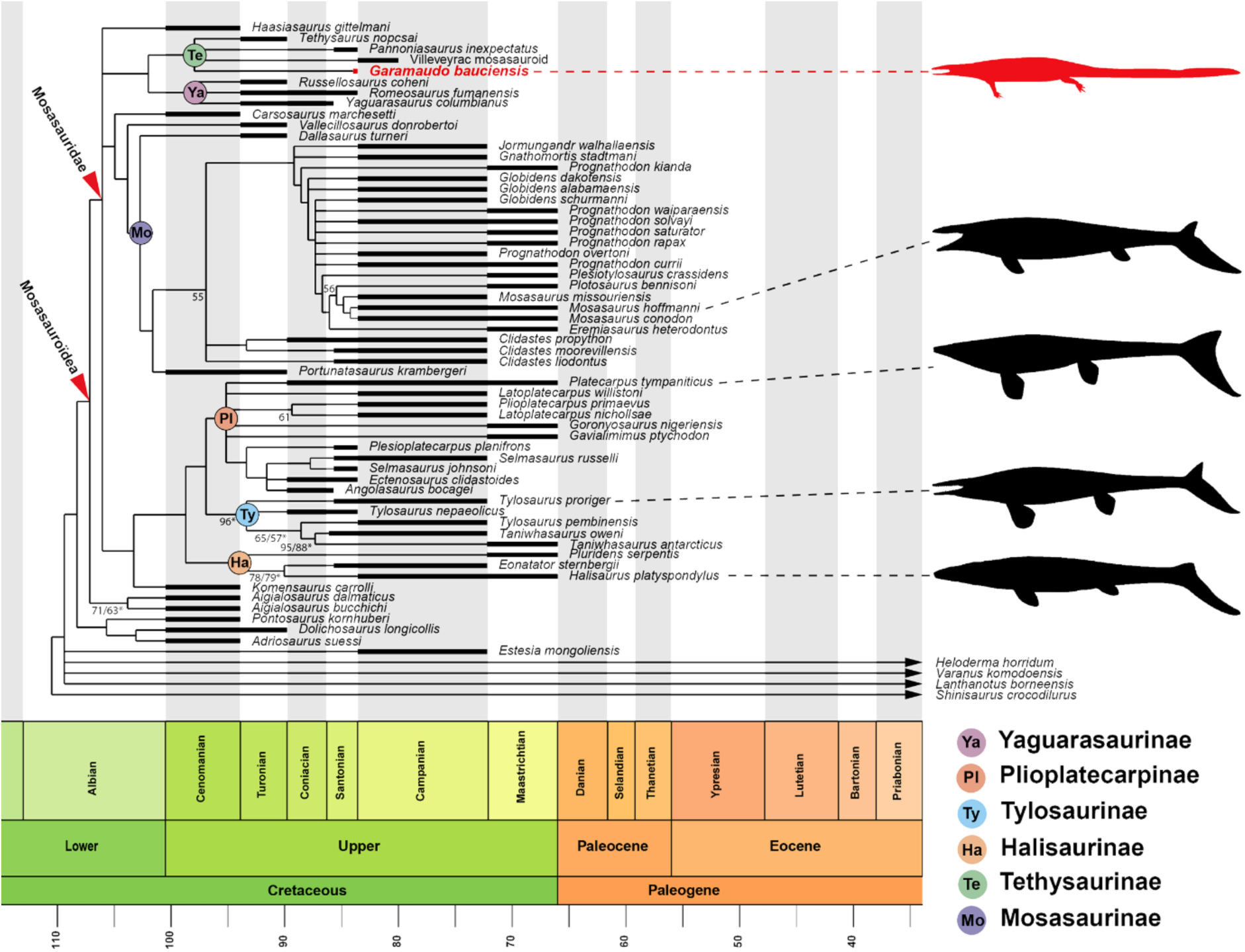
Time-scaled phylogeny of Mosasauroidea showing the position of *Garamaudo bauciensis* gen. et sp. nov. Based on the strict consensus tree obtained from the unweighthed analysis. International Chronostratigraphic Chart after Cohen *et al*. (2013). Temporal occurrences of the extinct taxa are represented as thick black bars whereas extant taxa are indicated with arrows. Node values correspond to bootstrap and jackknife (asterisk) indices, only values equal or superior to 50 percent are indicated. Mosasaurid silhouettes from PhyloPic (http://phylopic.org): *Pannoniasaurus* (serving as model for *Garamaudo*) by Makádi, Caldwell & Ősi (CC BY 4.0); *Platecarpus* by Lindgren, Caldwell, Konishi & Chiappe (CC BY-SA 3.0); *Tylosaurus* by Hartman (CC BY 3.0); *Mosasaurus* by Robinson (CC BY-NC 3.0); Silhouettes of Halisaurinae and *Garamaudo* (modified from *Pannoniasaurus*) designed by F.-L. Pelissier.

*Garamaudo* is recovered as a mosasaurid, more specifically as a member of Tethysaurinae (*sensu* Makádi *et al*., 2012).

Tethysaurinae is recovered as the sister-group of the Yaguarasaurinae (*sensu* Palci *et al*., 2013), both forming an unnamed clade that remains unresolved at the base of Mosasauridae. The mosasaurid subfamilies Halisaurinae, Plioplatecarpinae, Tylosaurinae, and Mosasaurinae are recovered as in previous analyses (e.g., Zietlow *et al*., 2023).

Node support values are low (<50%) for Tethysaurinae (1), Yaguarasaurinae (1) and the unamed clade (2), indicating limited robustness of these groupings in the present analysis.

The unnamed clade including Tethysaurinae and Yaguarasaurinae is supported by 10 unambiguous synapomorphies: parietal table elongate, triangular to subrectangular; narrow postorbitofrontal; postorbitofrontal transverse dorsal ridge present; quadrate suprastapedial process without dorsal constriction; basioccipital canal present; dentary medial parapet elevated and straplike; surangular-articular suture at the middle of the glenoid fossa in lateral view; synapophyses extending far below the ventral margin of the centrum; type 2 vascularization in the basisphenoid; type 2 vascularization in the basioccipital, with the posteromedial canal entering the basisphenoid between the carotid artery path and the abducens nerve exit.

Yaguarasaurinae *sensu* Palci *et al*. (2013) includes *Yaguarasaurus*, *Russellosaurus* and *Romeosaurus* in a polytomy. They are supported by four unambiguous synapomorphies: frontal not invaded by the posterior end of the nares; long quadrate suprastapedial process that extends below midheight; shallow quadrate alar concavity; ectopterygoid process of the pterygoid offset anterolaterally and bearing longitudinal grooves and ridges.

Tethysaurinae *sensu* Makádi *et al*. (2012) includes *Tethysaurus*, *Pannoniasaurus*, *Garamaudo* and the Villeveyrac mosasauroid, also in a polytomy. The clade is supported by seven unambiguous synapomorphies: premaxilla arcuate anteriorly; low median frontal dorsal keel; frontal with discrete ornamentations; 17–19 dentary teeth; humerus ectepicondyle present; humerus entepicondyle present; short retroarticular process relative to glenoid.

Among tethysaurines, *Garamaudo bauciensis* bears one unambiguous apomorphy: condyle of posterior trunk vertebrae slightly compressed. However, the analysis fails to resolve its position relative to other tethysaurines. This poor resolution could result from the incompleteness of most of these basal taxa.

## Discussion

### Systematic attribution

Both the anatomical and microanatomical characters observed in the cranial, axial and appendicular skeleton of *Garamaudo* permit its reference to a pythonomorph squamate, and more specifically to a Mosasauroidea (*sensu* Bell & Polcyn, 2005; Caldwell & Palci, 2007).

*Garamaudo* differs greatly from hydropedal and hydropelvic mosasaurids (*sensu* Bell & Polcyn, 2005; Caldwell & Palci, 2007) that bear: a posterior mandibular unit (PMU) proportionally shorter, deeper and more diamond-shaped in lateral view, centra that are more cylindrical and less flat, a humerus that is much shorter and broader, an unfused pelvis with an anteriorly oriented ilium (e.g., Russell, 1967; DeBraga & Carroll, 1993; Zietlow *et al*., 2023), and a much lighter microanatomical structure made of a tight trabecular network (Houssaye & Bardet, 2012).

It differs also from the basal plesiopedal / hydropelvic mosasaurine *Dallasaurus* that, though exhibiting a comparably long and slender humerus, possesses more cylindrical vertebral centra, a slightly shorter humerus with more expanded epiphyses, a pelvis with a loose acetabular articulation and an anteriorly oriented ilium (hydropelvic condition), vertebrae with a much lower compactness, and ribs with an open medullary cavity (Bell & Polcyn, 2005; Houssaye & Bardet, 2012).

Conversely, most of its plesiomorphic characteristics, such as a long and slender PMU, a long and slender humerus with epiphyses almost perpendicularly oriented from one another, and a pelvis with fused acetabulum articulation and posteriorly oriented ilium, are reminiscent of basal mosasauroids, including plesiopedal and plesiopelvic “aigialosaurids” and tethysaurines, as well as yaguarasaurines, whose appendicular condition remains mostly unknown (e.g., Palci *et al*., 2013; Zietlow *et al*., 2023). In the details, however, *Garamaudo* differs from all these taxa as follows.

The pythonomorph *Mesoleptos* (Lee & Scanlon, 2002) shares with *Garamaudo* a comparable rib morphology, indicating a probable laterally compressed body, and a humerus with a semi-lunar proximal epiphysis and a long and slender shaft although remaining small. It however differs from *Garamaudo* in a slender and twisted humeral shaft and a stronger vertebral osteosclerosis (Houssaye, 2013a).

*Aigialosaurus* (*A*. *dalmaticus* and *A*. *bucchichi*) is comparable to *Garamaudo* in its long and slender PMU, rib morphology, long and slender humerus, and posteriorly oriented ilium. However, *Aigialosaurus* differs in having an ilium that bears a posterior process that wide and robust along its length, and a large preacetabular process, as well as by possibly (since they are only slightly swollen) pachyostotic ribs (Dutchak & Caldwell, 2006, 2009).

*Komensaurus* is comparable to *Garamaudo* in its rib morphology and the posteriorly oriented ilium with short preacetabular process. However, it differs in having a pelvis exhibiting a loosely sutured acetabulum and a shorter, distally fan-shaped pubis (Caldwell & Palci, 2007).

*Carsosaurus* has a long and slender humerus and a posteriorly oriented ilium comparable to *Garamaudo*. However, it differs in having well-developed pectoral and deltoid crests of the humerus and by its long, convex ribs (Caldwell *et al*., 1995).

*Vallecilosaurus* bears a posteriorly oriented ilium, similarly as *Garamaudo*, but its pubis appears shorter (Smith & Buchy, 2008). Further comparisons are not possible due to the lack of detailed figures.

*Haasiasaurus* has a long and slender PMU, a long and slender humerus and a posteriorly oriented ilium, as *Garamaudo*. Its PMU, however, differs greatly in having a surangular that is very concave dorsally and bears a large foramen anterolaterally, a fan-shaped (as in plioplatecarpines), posteromedially oriented retroarticular process, and a glenoid fossa made equally by the surangular and articular (Polcyn *et al*., 1999). *Haasiasaurus* differs also from *Garamaudo* in having a pelvis with loose acetabular articulation and vertebrae exhibiting more extensive osteosclerosis and some pachyostosis (Polcyn et al., 1999; Houssaye, 2013a).

The yaguarasaurine *Yaguarasaurus* differs from *Garamaudo* in having a much deeper PMU (Páramo-Fonseca, 1994).

The yaguarasaurine *Russellosaurus* has a PMU comparable in general shape to *Garamaudo*, notably in its long and slender morphology, elongated slender coronoid articulation, and glenoid fossa made mostly by the articular, the surangular participating only anteriorly (Polcyn & Bell, 2005). However, it differs in having a retroarticular process that is ovoid and posteroventrally oriented, and a shorter surangular with a concave dorsal margin (between the glenoid and coronoid) (in *Garamaudo*, it is longer and straight).

The yaguarasaurine *Romeosaurus* exhibits a PMU that is long and slender, as in *Garamaudo*, with a surangular bearing a coronoid suture long and slender and a lateral articular suture ‘V’-shaped, as well as a short retroarticular process. It differs by its possession of vertebral hypapophyses with rounded articular surfaces for the hypocentrum, ribs with rounded heads, and a much shorter and wider humerus (Palci *et al*., 2013).

The tethysaurine *Tethysaurus* possesses a PMU and a posteriorly oriented ilium comparable in shape to *Garamaudo*. However, it differs in having a triangular, longer and obliquely oriented retroarticular process, a pelvis bearing loose acetabular articulation and a shorter, distally fan-shaped pubis (Bardet *et al*., 2003). *Tethysaurus* shows no osteosclerosis, either (Houssaye & Bardet, 2012).

The tethysaurine *Pannoniasaurus*, a coeval freshwater taxon from Hungary, is similar to *Garamaudo* in having a long and slender PMU, prezygapophyses that project strongly anteriorly, extending well beyond the anterior border of the cotyle, oval rib heads, a relatively flat proximal humeral epiphysis, and an ilium with a long and slender, posteriorly oriented, posterior process (Makádi *et al*., 2012). However, *Pannoniasaurus* differs in having a longer, obliquely oriented retroarticular process bearing a large foramen, rounded cervical hypapophyses, no zygosphenal foramen, and a pelvis with probably loose acetabular articulation.

The Villeveyrac mosasauroid, a possible tethysaurine also from almost coeval freshwater environments of southern France (Garcia *et al*., 2015), has similar vertebrae to those of *Garamaudo.* Unfortunately, no more material is available for comparisons and the microanatomy is unknown.

In conclusion, *Garamaudo bauciensis* can be referred to the Tethysaurinae (*sensu* Makádi *et al*., 2012) on the basis of the following characters: mandibular glenoid formed mainly by the articular, cervical synapophyses extending below the ventral border of the centrum, ilium with long and slender posterior process and spoon-shaped preacetabular process overlapping the pubis.

Though retaining several plesiomorphic characters, *Garamaudo* differs however from all basal mosasauroids as shown above, and is unique among tethysaurines, being characterized by the following autapomorphies (detailed in the diagnosis): PMU strongly curved inward in dorsal/ventral view; surangular with a medial horizontal triangular plateau anterior to the glenoid fossa, followed by a straight adductor fossa, open along its length; retroarticular process extremely short and semi-lunar, oriented obliquely at about 45°, extending anteromedially, and ending as an anteriorly oriented hook; osteosclerotic vertebrae; ribcage dorsoventrally deep and laterally compressed; humerus highly osteosclerotic, with a long and slender diaphysis, possibly reduced; pubis with a very long and slender anterior process, oriented parallel to the long axis of the body.

### Palaeobiology

On the basis of the preserved bones of *Garamaudo*, assumed to belong to adult specimens (see ‘General preservation’), its size is estimated to about 2.5 m long. Notably, its PMU and vertebrae are comparable in size to those of *Tethysaurus*, estimated to be less than 3.0 m long (Bardet *et al*., 2003). The Villeveyrac mosasauroid, a possible tethysaurine, was also estimated to be around 3 m long (Garcia *et al*., 2015). Only *Pannoniasaurus* appears larger, having been estimated to have been a medium-sized tethysaurine up to 6 m long (Makádi *et al*., 2012).

Palci *et al*. (2013) mentioned that, as basal mosasauroids (“aigialosaurids”), *Tethysaurus* and *Pannoniasaurus* retain the plesiopedal and plesiopelvic conditions. This also applies to *Garamaudo* and to the Villeveyrac mosasauroid (Garcia *et al*., 2015), given the morphology of their preserved limb and girdle bones.

Vascularization indicates that the growth rate and, by extrapolation, metabolism of *Garamaudo* were probably similar to those of some pythonomorphs (*Pachyvaranus*, “pythonomorph” from Touraine, *Carentonosaurus*) (Buffrénil *et al*., 2008; Houssaye *et al*., 2008, 2010), which share a similar vascularization pattern. *Garamaudo* was thus probably more active than modern lizards (which have limited or absent vascularization) but less so compared to the supposedly gigantothermic mosasaurids (exhibiting higher vascularization) (see Houssaye, 2013b).

There is clear osteosclerosis in all studied postcranial bones of *Garamaudo*. This implies a hydrostatic control of buoyancy and body trim (Houssaye, 2009), which suggests an animal capable of diving to shallow depths with possibly limited acceleration and maneuverability, indicative of ambush predation. This osteosclerosis also appears incompatible with terrestrial locomotion, because of the high risk of fracture, despite the otherwise possible plesiopedal and plesiopelvic conditions of its appendicular skeleton. The individuals that comprise the hypodigm of *Garamaudo* probably lived in the local freshwater environment where their bones were preserved (see below). The plesiopedal and plesiopelvic conditions of the animal also suggest that its mode of swimming was mainly anguilliform (e.g., Lindgren *et al*., 2011).

It is worth noting that the long and nearly straight ribs indicate a laterally compressed body that is deeper than wide. The proportionally small humerus suggests a reduced forelimb. The only element of *Garamaudo* permitting comparison of humeral length to body size is the PMU. For this purpose, a ratio of humerus length / PMU length was calculated. Unfortunately, very few ophidiomorphs and basal mosasauroids preserve both elements. The ratio of *Garamaudo* (0.55) is comparable to that of *Haasiasaurus* (0.53), but remains much lower compared to other basal mosasauroids like *Aigialosaurus dalmaticus* (0.63) and *A. bucchichi* (0.87). It is rather similar to that of ophiodomorphs like *Pontosaurus kornhuberi* (0.59) and higher than that of *Pontosaurus lesinensis* (0.37). This makes the proportions of the humerus of *Garamaudo* rather consistent with those observed in these other plesiopedal taxa, although the size difference could simply be due to the fact that the humerus belongs to a smaller individual than the holotype.

### Palaeoenvironment

The Bouc-Bel-Air - Sousquières locality (BBAS) represents a freshwater ecosystem, probably a moderately eutrophic, carbonate-producing lake periodically influenced by flooding and shallowing events (see Geology), within the continental deposits of the Aix-en-Provence Basin (Babinot & Durand, 1980; Tortosa & Leleu, 2021). Such rapid flooding episodes could explain the state of preservation of the holotype specimen of *Garimaudo*, which includes broken bones having however well-preserved surfaces. The vertebrate assemblage of BBAS includes the ambush actinopterygian predator *Atractosteus* cf. *africanus* (∼2 m; Cavin *et al*., 1996, 2020; “lepisosteidae indet.” in Tortosa, 2014), the benthic coelacanth *Axelrodichthys megadromos* (∼1 m; Cavin *et al*., 2020), bothremydid turtle cf. *Polysternon* sp. (∼60 cm), small indeterminate crocodylomorphs, and *Garamaudo*, a mid-sized mosasauroid (∼2.5 m).

As in the Lower Campanian of Villeveyrac locality (Garcia *et al*., 2015), no large crocodylomorphs are known from BBAS or equivalent Valdonnian–lower Fuvelian localities in Provence (Cavin *et al*., 1996; Tortosa, 2014; Cavin *et al*., 2020). *Garamaudo* therefore ranks among the largest predatory vertebrates in the assemblage, together with *Atractosteus*. Although its precise diet remains unknown, *Garamaudo* may have fed on small fishes, reptiles and/or amphibians, either by active predation and/or scavenging (Figure 11).

**Figure 11.**
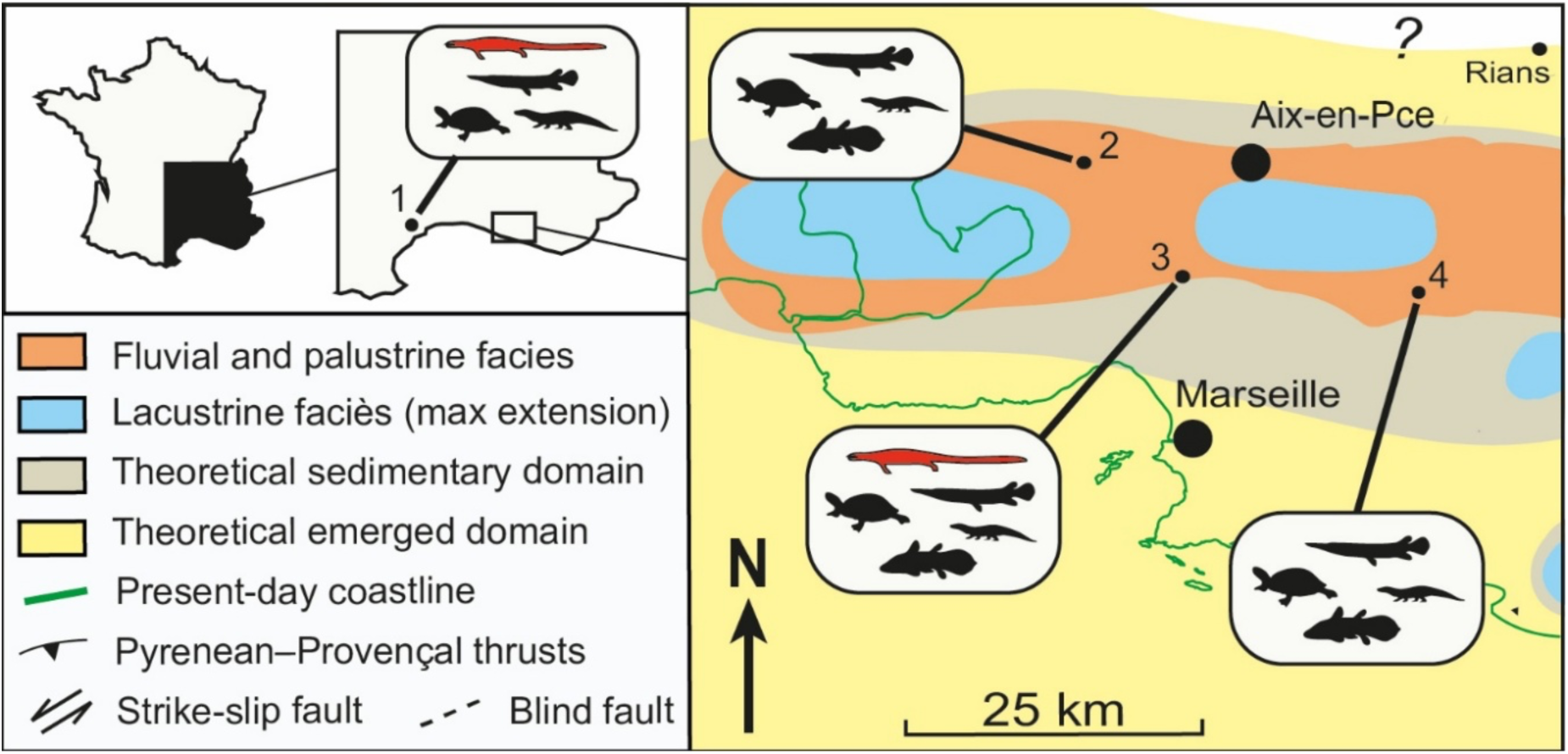
Paleogeographic reconstruction maps of Provence continental deposits during the latest Santonian (Valdonnian facies), after Tortosa & Leleu (2021), with indications of the main vertebrate outcrops and their faunal content (larger vertebrates only). **1**, Villeveyrac (Languedoc, early Campanian, Garcia *et al*., 2015); **2**, Ventabren (Cavin *et al*., 2020); **3**, Bouc-Bel-Air (Cavin *et al*., 2020; this study); **4**, Belcodène (Tortosa, 2014). See Figure 2 for silhouette legend (*Garamaudo bauciensis* gen. et sp. nov. in red).

In younger Fuvelian lignite levels, the emergence of medium-sized crocodylomorphs (*Massaliasuchus* and *Allodaposuchus*-like forms) is associated with a marked shift in the fish fauna, which becomes dominated by smaller actinopterygians such as *Lepisosteus* sp. (Martin & Buffetaut, 2008; Tortosa, 2014). At the same time, mosasauroids are no longer recorded in these deposits, whereas other aquatic predators, including hybodontid sharks, appear (Tortosa, 2014; Valentin *et al*., 2025).

The trophic structure of BBAS parallels that of the Hungarian site of Iharkút, where *Pannoniasaurus*, a tethysaurine mosasaurid closed to *Garamaudo* (see Phylogeny), coexisted with bothremydid turtles, small crocodiles, and gar-like fishes within a freshwater meandering system (Makádi *et al*., 2012). The BBAS locality thus represents the second westernmost known equivalent of these continental freshwater ecosystems in Europe during the Santonian, showing that mosasauroids could successfully share top-predator niches in lacustrine-palustrine environments.

### Palaeoecology

Modern squamates are known to occupy a wide range of habitats, from purely marine to continental ones, the latter spanning estuarine, mangrove, brackish, and fresh-water environments (Carrillo *et al*., 2024; Neill, 1958). Single species can also exploit various environments alternatively, switching from one to another for either food access, reproductive purposes, or to escape predators, as seen in some Southeast Asian island species of *Varanus* (Pianka & King, 2004). Similarly, *Crocodilus porosus*, the only “true” marine living crocodile, is also encountered in estuarine, brackish, and swampy habitats and can be found far inland in freshwater environments (Webb *et al*., 2010). Extant marine reptiles are thus able to occupy an extended range of aquatic environments, attesting to their adaptability, especially in terms of salinity levels.

Although the oldest (Cenomanian-Turonian) mosasauroids (“aigialosaurids”) exhibit limited morphological adaptations to an aquatic lifestyle, all have been found in coastal marine environments (Palci *et al*., 2008), suggesting that the clade rapidly evolved from continental (terrestrial or freshwater) squamates into marine forms. Turonian mosasaurids, including tethysaurines (*Tethysaurus*), yaguarasaurines *(Yaguarasaurus*, *Russellosaurus*, *Romeosaurus*) and early mosasaurines (*Dallasaurus*), also occurred in marine or estuarine environments (e.g., Polcyn *et al*., 2025), illustrating that, at that time, they occupied various ecological niches there. From the Coniacian–Santonian onward, mosasaurids underwent a progressive shift toward more open-sea conditions until their extinction (e.g., Bardet *et al*., 2014; Polcyn *et al*., 2014, 2025). Mosasauroids have thus generally been considered marine taxa. However, this has been challenged by recent discoveries of continental tethysaurine mosasauroid remains (Makadi *et al*., 2012; Garcia *et al*., 2015).

Among tethysaurines, whereas the Turonian *Tethysaurus* occupied a clearly marine environment (see Bardet *et al*., 2003), the Santonian *Pannoniasaurus* and early Campanian Villeveyrac mosasauroid lived in predominantly freshwater continental ecosystems (Makádi *et al*., 2012; Garcia *et al*., 2015). Isolated tethysaurine teeth were also reported from Turonian and Coniacian freshwater ecosystems of Austria (Ősi *et al*., 2019, 2021). The attribution of *Garamaudo* to tethysaurines thus illustrates an additional freshwater occurrence for this clade. Except for a *Plioplatecarpus* found in estuarine Maastrichtian deposits of Canada (Holmes *et al*., 1999), this mosasaurid subfamily is the only one currently known to have evolved multiple species autochtonous in freshwater environments. Given the paleogeography of the Late Cretaceous European Archipelago (Csiki-Sava *et al*., 2015), if these multiple occurrences resulted from a single evolutionary transition from marine to freshwater environments, it would imply subsequent migrations into various remote freshwater environments through marine routes. This rather suggests several independent conquests of freshwater habitats.

Although the occurrence of these taxa in freshwater deposits could represent excursions from marine environments, the absence of their remains in marine deposits, and the presence of three species so-far known in such ecological contexts, reinforce the likelihood of their freshwater habits. The ecological plasticity observed in some modern aquatic reptiles could explain the establishment of freshwater tethysaurines across the Late Cretaceous European archipelago, allowing them to exploit a mosaic of new ecological niches, facilitated by the absence of large crocodylomorphs, the usual apex-predators of these continental ecosystems (Kocsis *et al*., 2009; Makádi *et al*., 2012; see above). Freshwater tethysaurines may also have found refuges inland, facilitating ecological niche partitioning and thereby reducing direct competition with large (> 5 m) coeval marine mosasaurids that became increasingly abundant at this time, including in the European archipelago (e.g., Plasse *et al*., 2024; Polcyn *et al*., 2025).

As already mentioned, all tethysaurines known up to now retain the plesiopedal and plesiopelvic conditions. Except *Tethysaurus*, all have been found in continental freshwater environments, and, except *Pannoniasaurus*, all are reduced in size, being no more than 3 m long. The freshwater environment might have supported the retention of globally small sizes and plesiomorphic limbs and girdles among this basal mosasaurid clade.

## Conclusion

New mosasauroid remains from the Santonian of Provence, southeastern France are attributed to a new tethysaurine mosasaur: *Garamaudo bauciensis*. The Bouc-Bel-Air - Soucquières (BBAS) locality corresponds to a carbonate-producing freshwater lake and represents the second-most westerly Late Cretaceous continental freshwater ecosystem known in Europe. *Garamaudo* is about 2.5 m long, with a plesiopedal and plesiopelvic morphology, and displays osteosclerosis in its vertebrae, ribs, and humerus. The sedimentological, anatomical, and microanatomical data, as well as its estimated intermediate growth rate and metabolism compared to other mosasaurids, are consistent with an anguilliform swimmer foraging underwater at shallow depths.

The assignment of this new freshwater mosasaurid to the tethysaurines highlights an additional occurrence of a freshwater species in this clade and the fact that, based on current knowledge, it is only within the Tethysaurinae that freshwater forms evolved among mosasaurids. During the same period, hydropedal and hydropelvic mosasaurs with a higher growth rate and spongious bones, indicating an active carangiform swimming mode, evolved and radiated in open marine environments, spreading throughout the world. This new discovery therefore contributes to a better understanding of a more diverse and complex evolution and history of post-Turonian mosasaurids, including more extensive niche partitioning than previously realized.

## Supporting information

Appendix 1 (Phylogeny)

Matrix

## Acknowledgments

We gratefully acknowledge the *Département des Bouches-du-Rhône* and J.-M. Perrin, its elected representative for the promotion of Provence’s paleontological and archaeological heritage, for their continuous support, excavation authorization, and administrative assistance. We are also indebted to the *Direction de l’Environnement, des Grands Projets et de la Recherche* (E. Mangion, H. Souan, M. Bourrelly) and to the *Direction des Routes et des Ports* (F. Cauvin, C. Maréchal, B. Ott, A. Hémery, P. Abignoli) for their help in the field and logistical coordination. We further thank the staff of the *Muséum d’Histoire Naturelle d’Aix-en-Provence* (S. Berton, M. Desparoir, N. Vialle). We acknowledge the MRI platform member of the national infrastructure France-BioImaging supported by the French National Research Agency (ANR-10-INBS-04, «Investments for the future»), the labex CEMEB (ANR-10-LABX-0004) and NUMEV (ANR-10-LABX-0020) and notably R. Lebrun for facilitating access to and use of the µCT. We also warmly thank V. Fischer (Liège University, Belgium) for helpul discussions on the Bremer support indices, as well as P. Loubry (CR2P, Paris) and A. Lethiers (ISTeP / CR2P, Paris) for respectively the photographs and drawings / design of the *Garamaudo* figures. Finally, we thank an anonymous referee and T. Konishi (University of Cincinnati, USA) for their comments and J. Mallon for his editorial work, which allowed to greatly improve the quality of our manuscript.

## Author contributions

Conceptualization: NB, AH, TT; Fieldwork and preliminary investigation: DY, ET, TT; Data curation: DY, TT; Data acquisition: AH; Formal analysis: NB, AH, FLP, TT; Writing – original draft: NB, AH, FLP, TT; Final validation by all authors.

## Data, scripts, code, and supplementary information availability

Data, script, and supplementary information are available online: https://doi.org/10.64898/2025.12.11.693649.

