## Appendix 1 (Phylogeny) for "*Garamaudo bauciensis*, a new freshwater Mosasauridae (Reptilia, Squamata) from the Santonian (Late Cretaceous) of Provence, southeastern France"

**Taxa**

**Added taxa:**

***Garamaudo bauciensis*, nov. gen. nov sp., this work.**

Villeveyrac mosasauroid, scored from the description by Garcia *et al.* (2015).

*Vallecillosaurus donrobertoi*, scored from the description by Smith & Buchy (2008) and matrix by Mekarski (2017).

*Haasiasaurus gittelmani*, scored from the description by Polcyn *et al.* (1999) and matrix by Mekarski (2017).

*Carsosaurus marchesetti*, scored from the description by Kornhuber (1893), Caldwell *et al.* (1995) and matrix by Mekarski (2017).

*Portunatasaurus krambergeri*, scored from the description by Mekarski *et al.* (2019) and matrix by Mekarski (2017).

**Modified taxa:**

*Shinisaurus crocodilurus* modified from Augusta *et al.* (2022). Character 43 (1 🡪 0).

*Heloderma horridum* modified from Augusta *et al.* (2022). Character 43 (1 🡪 0).

*Lanthanotus borneensis* modified from Augusta *et al.* (2022). Character 43 (1 🡪 0).

*Varanus komodoensis* modified using observation of the 3D model TNHC.Herpetology.95803 (Gray, 2024). Character 43 (1 🡪 0).

*Adriosaurus suessi* modified from Augusta *et al.* (2022). Character 43 (? 🡪 0).

*Pontosaurus kornhuberi* modified from Augusta *et al.* (2022). Character 43 (? 🡪 0).

*Agialosaurus dalmaticus* modified from Augusta *et al.* (2022) and Dutchak & Caldwell (2009). Character 25 (0 🡪1), character 85 (1 🡪 0).

*Aigialosaurus* (= *Opetiosaurus*) *bucchichi* modified from Augusta *et al.* (2022) and Dutchak & Caldwell (2009). Character 25 (0 🡪 1), character 43 (0 🡪 1), character 85 (1 🡪 0), character 86 (? 🡪 0).

*Komensaurus carrolli* modified from Augusta *et al.* (2022). Character 43 (0 🡪 1).

*Pannoniasaurus inexpectatus* modified from Makádi *et al.* (2012). Character 4 (? 🡪 0), character 28 (? 🡪 1), character 37 (0 🡪) 1; character 43 (0 🡪 1), character 47 (2 🡪 0), character 53(0 🡪 ?), character 62 (? 🡪 1), character 63 (? 🡪 0), character 69 (1 🡪 0), character 70 (1 🡪 ?), character 72 (0 🡪 1), character 78 (1 🡪 0), character 79 (1 🡪 0), character 84 (0 🡪 ?), character 98 (0 🡪 ?), character 99 (1 🡪 ?), character 102 (? 🡪 0), character 103 (? 🡪 0), character 104 (? 🡪 1), character 106 (0 🡪 1), character 107 (0 🡪 1).

*Tethysaurus nopcsai* modified from Bardet *et al.* (2003), Makádi *et al.* (2012), Houssaye & Bardet (2012) and Augusta *et al.* (2022). Character 10 (1 🡪 0), character 13 (? 🡪 0), character 15 (2 🡪 3, character 16 (? 🡪 0), character 43 (0 🡪 1), character 78 (1 🡪 0), character 79 (1 🡪 0), character 94 (1 🡪 0), character 106 (0 🡪 1), character 107 (0 🡪 1).

*Romeosaurus fumanensis* modified from Palci *et al.* (2013). Character 7 (? 🡪 0), character 8 (? 🡪 2).

*Pluridens serpentis* modified from Longrich *et al.* (2021). Character 15 (? 🡪 3).

*Eonatator sternbergi* modified from Augusta *et al.* (2022). Character 43 (0 🡪 1).

*Halisaurus platyspondylus* modified from Bell (1997). Character 43 (0 🡪 1).

**Characters**

**Modified character:**

40. Quadrate suprastapedial process fusion: absent (0); present with a pillar-like process in some case (1); present with the base of quadrate (2). Based on Palci *et al.* (2021).

**Added characters:**

125. Caudal vertebrae, depth: Regular, no sculling organ, ratio being 1,4 or less (0); weakly developed sculling organ, ratios varying from 1,5 to 2,5 (1); very deep, forming a strong sculling organ, ratio being 2.6 or higher (2). Based on Augusta *et al.* (2022).

126. Posteromedial canal enters basisphenoid between the carotid artery path and abducens nerve exit: absent (0); present (1). Based on character 98 of Polcyn *et al.* (2023).

127. Vascularization of the basisphenoid and basioccipital: Type 1, no accessory anterolateral foramina or internal canal in the basisphenoid or basioccipital (0); Type 2, posteromedial canal in the basisphenoid present, forming a canal or vestibule in the basisphenoid body, exiting ventrally and does not pass into basisphenoid anteriorly (1); Type 3, posteromedial canal in the basisphenoid present, forming a canal or vestibule in the basisphenoid body, and passes anteriorly into basisphenoid (2). Note: this character is the fusion of characters 99 and 100 of Polcyn *et al.* (2023); states were coded as 0 (inferred) when not observed in Halisaurinae, Mosasaurinae, and Tylosaurinae, following Polcyn *et al.* (2023); states were coded as “?” in dolichosaurids and basal branching mosasauroids, also according to Polcyn *et al*. (2023).

128. Dorsal surface of the frontal bears discrete ornamentations: absent (0); present (1).

129. Retroarticular process length relative to glenoid fossa: elongated (0); short (1); extremely short (2). Note: the length is qualitative here and based on personal observations and descriptions using bibliography. “Elongated” refers to a long, slender retroarticular process, longer than the glenoid fossa (as in outgroups such as varanids); “Short” refers to a curved retroarticular process approximately equal in length to the glenoid fossa (as in most mosasauroids); “Extremely short” refers to a curved retroarticular process distinctly shorter than the glenoid fossa (as in *Garamaudo*).

**Outgroup list**

| Species | Family | Superfamily | Reference |
| --- | --- | --- | --- |
| *Shinisaurus crocodilurus* | Shinisauridae | — | Ahl (1930) |
| *Lanthanotus borneensis* | Lanthanotidae | Varanoidea | Augusta *et al.* (2022) |
| *Varanus komodoensis* | Varanidae | Varanoidea | Augusta *et al.* (2022) |
| *Heloderma horridum* | Helodermatidae | Varanoidea | Augusta *et al.* (2022) |
| *Estesia mongoliensis* | — | Varanoidea | Norell *et al.* (1992) |
| *Adriosaurus suessi* | « Dolichosauridae » | — | Augusta *et al.* (2022) |
| *Dolichosaurus longicollis* | « Dolichosauridae » | — | Augusta *et al.* (2022) |
| *Pontosaurus kornhuberi* | « Dolichosauridae » | — | Augusta *et al.* (2022) |

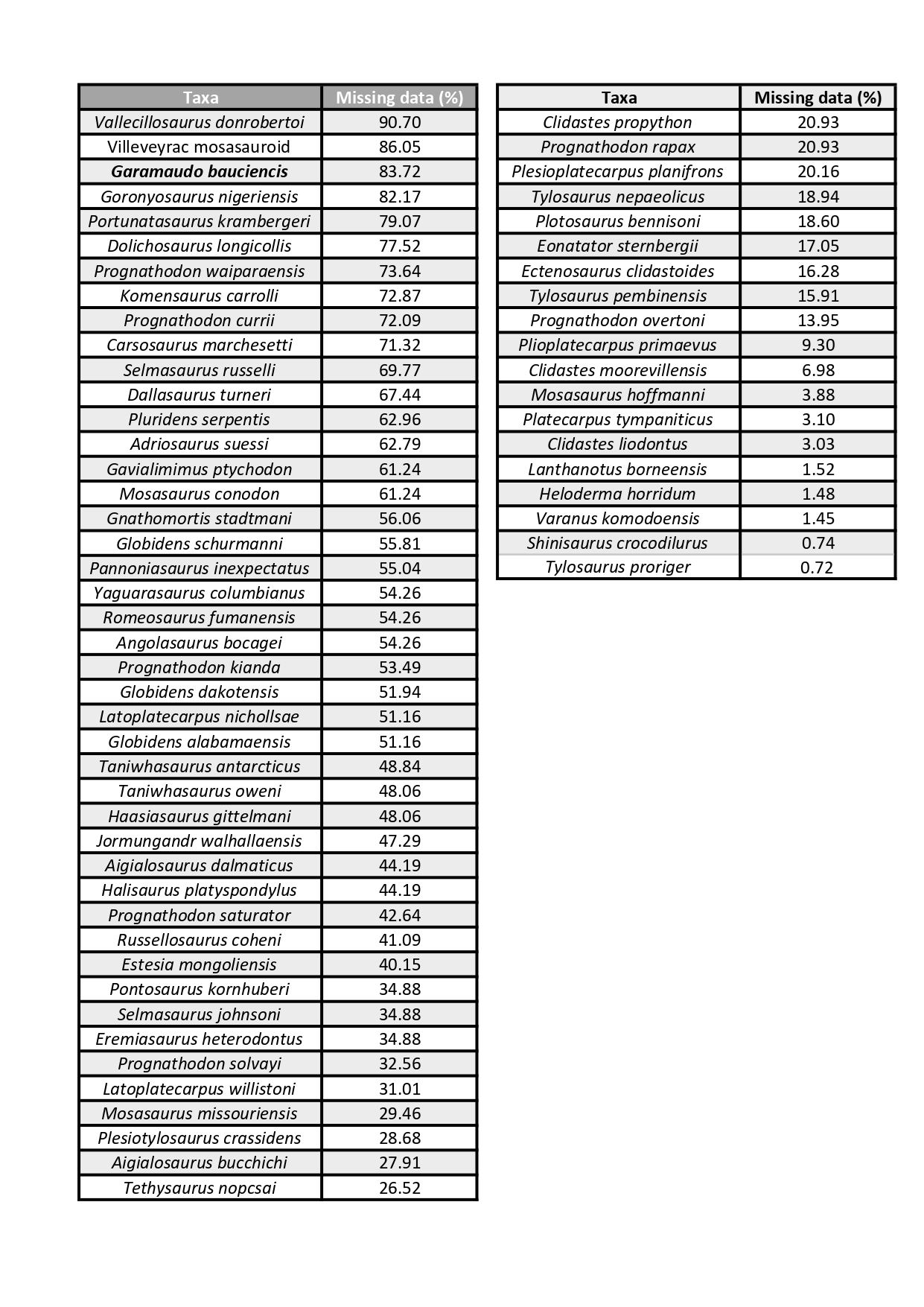
**Missing data ( ?) percentage for each taxon**
