## Supplementary material for "*Garamaudo bauciensis*, a new freshwater Mosasauridae (Reptilia, Squamata) from the Santonian (Late Cretaceous) of Provence, southeastern France": Matrix

```
#NEXUS
[written Mon Dec 15 18:30:13 CET 2025 by Mesquite  version 3.81 (build 955) at FL_PALEO/192.168.1.13]

BEGIN TAXA;
	TITLE Taxa;
	DIMENSIONS NTAX=63;
	TAXLABELS
		'Shinisaurus_crocodilurus' 'Lanthanotus_borneensis' 'Varanus_komodoensis' 'Heloderma_horridum' 'Estesia_mongoliensis' 'Adriosaurus_suessi' 'Dolichosaurus_longicollis' 'Pontosaurus_kornhuberi' 'Aigialosaurus_bucchichi' 'Aigialosaurus_dalmaticus' 'Komensaurus_carrolli' Portunatasaurus_krambergeri 'Halisaurus_platyspondylus' 'Eonatator_sternbergii' 'Pluridens_serpentis' 'Yaguarasaurus_columbianus' 'Romeosaurus_fumanensis' 'Russellosaurus_coheni' Garamaudo_beauciensis 'Mosasauroid_of_Villeveyrac' 'Pannoniasaurus_inexpectatus' 'Tethysaurus_nopcsai' 'Taniwhasaurus_antarcticus' 'Taniwhasaurus_oweni' 'Tylosaurus_nepaeolicus' 'Tylosaurus_pembinensis' 'Tylosaurus_proriger' 'Angolasaurus_bocagei' 'Ectenosaurus_clidastoides' 'Gavialimimus_ptychodon' 'Goronyosaurus_nigeriensis' 'Latoplatecarpus_nichollsae' 'Latoplatecarpus_willistoni' 'Platecarpus_tympaniticus' 'Plesioplatecarpus_planifrons' 'Plioplatecarpus_primaevus' 'Selmasaurus_johnsoni' 'Selmasaurus_russelli' 'Dallasaurus_turneri' 'Clidastes_liodontus' 'Clidastes_moorevillensis' 'Clidastes_propython' 'Eremiasaurus_heterodontus' 'Gnathomortis_stadtmani' 'Mosasaurus_conodon' 'Mosasaurus_hoffmanni' 'Mosasaurus_missouriensis' 'Plesiotylosaurus_crassidens' 'Plotosaurus_bennisoni' 'Prognathodon_currii' 'Prognathodon_kianda' 'Prognathodon_overtoni' 'Prognathodon_rapax' 'Prognathodon_saturator' 'Prognathodon_solvayi' 'Prognathodon_waiparaensis' 'Globidens_alabamaensis' 'Globidens_dakotensis' 'Globidens_schurmanni' 'Jormungandr_walhallaensis' Vallecillosaurus_donrobertoi Haasiasaurus_gittelmani Carsosaurus_marchesetti
	;

END;


BEGIN CHARACTERS;
	TITLE  Character_Matrix;
	DIMENSIONS  NCHAR=129;
	FORMAT DATATYPE = STANDARD RESPECTCASE GAP = - MISSING = ? SYMBOLS = "  0 1 2 3 4 5";
	CHARSTATELABELS
		1 Pmx_predental_rostrum /  absent 'short, obtuse' distinctly_protruding 'large, inflated',
		2 Pmx_shape /  arcuate acute,
		3 Pmx_internarial_bar_width /  narrow wide,
		4 Pmx_internarial_bar_base_shape /  triangular rectangular 'T-shaped',
		5 Pmx_internarial_bar_dorsal_keel /  absent present,
		6 Pmx_internarial_bar_venter /  close_to_rostrum far_from_rostrum,
```

7 Frontal_shape_in_front_of_orbits /  sides_sinusoidal sides_straight,
8 Frontal_width /  'broad, short' intermediate 'long, narrow',
9 Frontal_narial_emargination /  absent present,
10 Frontal_midline_dorsal_keel /  absent low 'high, thin',
11 Frontal_ala_shape /  acuminate broad,
12 Frontal_olfactory_canal_embrasure /  absent present,
13 Frontal_posteroventral_midline /  'tabular boss anterior to frontal-parietal suture absent' boss_present,
14 'Frontal-parietal suture overlap orientation' / 'apposing; no overlap' oblique_median_frontal__&_parietal_ridges all_three_ridges_almost_horizontal,
15 Frontal_invasion_of_parietal_I / lateral_flange_of_frontal_posteriorly_extended medial_flange_of_frontal_posteriorly_extended both_extended suture_straight,
16 Frontal_medial_invasion_of_parietal_II /  'if present, posterior medial flange of frontal short' medial_frontal_flanges_long,
17 Parietal_length /  'dorsal surface short, epaxial musculature insertion posterior, between suspensorial rami only' 'dorsal surface long, epaxial musculature insertion posterior & dorsal',
18 Parietal_table_shape /  rectangular_to_trapezoidal triangular 'elongate, triangular to subrectangular, distinct mid- or parasagittal crest',
19 Parietal_foramen_size /  small large,
20 'Parietal foramen pos''n' /  near_center_of_parietal_table close_to_or_barely_touching_suture huge_foramen_invading_frontal,
21 Parietal_foramen_ventral_opening / level_with_main_ventral_surface surrounded_by_rounded_ridge,
22 Parietal_posterior_shelf / distinct_horizontal_shelf_projecting_posterior_to_horizontal_rami shelf_absent,
23 Parietal_susp_ramus_compression / greatest_width_vertical_or_oblique greatest_width_horizontal,
24 Parietal_union_with_supratemporal / parietal_overlaps_supratemporal_without_interdigitation forked_distal_ramus_sandwiches_proximal_end_of_supratemporal,
25 Prefrontal_supraorbital_process /  absent_or_very_small 'large, overhanging wing',
26 'Prefrontal-pof contact' /  absent prefrontal_overlapped_ventrally_by_postorbitofrontal prefrontal_overlapped_laterally_by_postorbitofrontal,
27 Pof_shape /  narrow wide,
28 Pof_transverse_dorsal_ridge /  absent present,
29 Mx_tooth_number /  '20-24' '17-19' '15-16' 14_or_less,
30 'Mx-pmx suture posterior terminus' / level_with_or_anterior_to_fourth_maxillary_tooth between_fourth_&_ninth_maxillary_teeth level_with_or_posterior_to_ninth_maxillary_tooth,
31 Mx_posterodorsal_extent /  posterodorsal_process_absent 'recurved wing of maxilla prevents prefrontal-naris contact' 'wing present but does not prevent prefrontal-naris contact',

32 Jugal_posteroventral_angle /  very_obtuse_to_curvilinear 'slightly obtuse, near 120 degrees' near_90_degrees,
33 Jugal_posteroventral_process /  absent present,
34 Ecp_contact_with_mx /  present absent,
35 Pterygoid_tooth_row_elevation /  absent teeth_arise_from_thin_pronounced_ridge,
36 Pterygoid_tooth_size / significantly_smaller_than_marginal_teeth 'large, approaching or meeting size of marginal teeth',
37 Q_suprastapedial_process_length /  short moderate long absent,
38 Q_suprastapedial_process_constriction / dorsal_constriction_present absent,
39 Q_suprastapedial_ridge /  'if present, ridge on ventromedial edge of process indistinct, straight, narrow' 'ridge wide, broadly rounded, curving downward',
40 Q_suprastapedial_process_fusion /  absent 'present with a pillar-like process in some case' present_with_the_base_of_Q,
41 Q_stapedial_pit_shape /  broadly_oval_to_circular narrow_oval elongate_with_constricted_middle,
42 Q_posteroventral_ascending_tympanic_rim_condition / ascending_ridge_small_or_absent 'high, elongate triangular crest' crest_extremely_produced_laterally,
43 Q_ala_thickness /  thin thick,
44 Q_conch /  deep shallow,
45 Basisphenoid_pterygoid_process_shape /  'narrow, articular surface facing anterolaterally' 'thinner, fan-shaped with posterior extension of articular surface causing more lateral orientation',
46 Q_ala_groove /  absent present_anterolaterally present_dorsally,
47 Q_median_ridge /  'single thin, high ridge' 'ridge low, rounded, diverging ventrally',
48 Q_anterior_ventral_condyle_modification /  absent anterodorsal_deflection_present,
49 Q_ventral_condyle /  'saddle-shaped, concave in posterior view' 'domed, convex',
50 Basioccipital_tubera_size /  short long,
51 Basioccipital_canal /  absent present_&_paired present_as_single_bilobate_cxanal,
52 Dentary_tooth_number /  more_than_20 '17-19' '15-16' 14 13 12_or_less,
53 D_anterior_projection /  absent short long,
54 D_medial_parapet /  at_base_of_tooth_roots 'elevated & strap-like, enclosing about half of height of tooth attachment in shallow canal' strap_equal_in_height_to_lateral_wall_of_bone medial_parapet_taller_than_lateral_wall_of_bone,
55 'Splenial-angular articulation shape' /  almost_circular laterally_compressed,
56 'Splenial-angular articular surface' /  smooth distinct_horizontal_tongues_&_grooves_present,
57 Coronoid_shape /  slight_dorsal_curvature 'very concave above, posterior wing fan-like',
58 Coronoid_posteromedial_process /  present_&_small absent,

59 Coronoid_medial_wing / does_not_reach_angular contacts_angular,
60 Coronoid_posterior_wing / medial_crescentic_pit_absent present,
61 Surangular_coronoid_buttress / 'low, thick, parallel to lower edge of jaw' 'high, thin, rapidly rising',
62 'Surangular-articular suture pos''n' / behind_glenoid_cotyle_in_lateral_view at_middle_of_cotyle anterior_to_cotyle,
63 'Surangular-articular lateral suture trace' / suture_curves_anteriorly suture_is_virtually_straight,
64 Articular_retroarticular_process_inflection / 'moderate inflection, less than 60 degrees' 'extreme inflection, almost 90 degrees',
65 Articular_retroarticular_process_innervation_foramina / absent one_to_three_large_foramina_present,
66 Tooth_surface_I / finely_striate_lingually lingual_striations_absent,
67 Tooth_surface_II / coarse_texture_absent coarse_texture_present,
68 Tooth_facets / absent present,
69 Tooth_fluting / absent present,
70 Tooth_inflation / absent present,
71 Tooth_carinae_I / absent present,
72 Tooth_carinae_serration / absent present,
73 Atlas_neural_arch / notch_in_anterior_margin notch_absent,
74 Atlas_synapophysis / extremely_reduced 'large, elongate',
75 Zygosphenes_and_zygantra / absent present,
76 Zygosphene_and_zygantra_number / present_on_many_vertabrae present_on_only_few_vertebrae,
77 Hypapophyses / last_hypapophysis_occurs_on_or_anterior_to_seventh_cervical on_eighth_cervical_or_posterior,
78 Synapophysis_height / 'tall, narrow on posterior cervicals & anterior dorsals' 'facets ovoid, shorter than centum height',
79 Synapophysis_length / 'on middle dorsals, not laterally elongate' distinctly_laterally_elongate,
80 Synapophysis_ventral_extension / 'minor, do not extend below ventral margin of centrum' extend_far_below_ventral_margin_of_centrum,
81 Vertebral_condyle_inclination / dorsal_vertebral_centra_inclined 'not inclined, vertical',
82 Vertebral_condyle_shape_I / 'condyles of anterior-most dorsals dorsoventrally depressed' essentially_equidimensional,
83 Vertebral_condyle_shape_II / condyles_of_posterior_dorsals_not_higher_than_wide slightly_compressed,
84 Vertebral_synapophysis_dorsal_ridge / absent sharp_ridge_&_anteriorly_precipitous_ridge_connecting_synapophysis_with_prezygapophysis,
85 Vertebral_length_proportions / cervicals_distinctly_shorter_than_longest_vertebrae cervicals_almost_equal_to_or_are_the_longest_vertebrae,

86 Presacral_vertebrae_number /  less_than_29 30_or_31 32_to_39 more_than_39,
87 Sacral_vertebrae_number /  two less_than_two,
88 Caudal_dorsal_expansion /  caudal_neural_spines_uniformly_shortened several_spines_dorsally_elongate_behind_middle_of_tail,
89 Haemal_arch_length /  about_equal_in_length_to_neural_arch_of_same_vertebra distinctly_longer_than_neural_arch_height,
90 Haemal_arch_articulation /  articulate_with_centra fused_to_centra,
91 Tail_curvature /  structural_downturn_absent present,
92 Body_proportions /  'head & trunk shorter than, or equal to, tail length' head_&_trunk_longer_than_tail,
93 'Scapula/coracoid size' /  both_bones_about_equal scapula_distinctly_smaller_than_coracoid,
94 Scapula_width /  anteroposterior_widening_absent 'distinct fan-shaped widening' extreme_widening,
95 Scapula_dorsal_convexity /  'if scapula widened, dorsal margin very convex' dorsal_margin_broadly_convex,
96 Scapula_posterior_emargination /  gently_concave deeply_concave,
97 'Scapula-coracoid suture' /  'unfused scapula-coracoid contact interdigitate anteriorly' apposing_surfaces_without_interdigitation,
98 Coracoid_neck_elongation /  neck_rapidly_tapering_to_broad_base neck_gradually_tapering_to_narrow_base neck_absent,
99 Coracoid_anterior_emargination /  present absent,
100 Humerus_length /  elongate shortened extremely_shortened distal_width_slightly_greater_than_length,
101 Humerus_postglenoid_process /  absent_or_very_small enlarged,
102 Humerus_glenoid_condyle /  'if present, gently domed, elongate, ovoid in proximal view' 'condyle saddle-shaped, subtriangular in proximal view & depressed' 'condyle highly domed or protuberant, short ovoid to almost round in proximal view',
103 Humerus_deltopectoral_crest /  undivided split_into_two_separate_insertional_areas,
104 Humerus_pectoral_crest /  located_anteriorly located_medially,
105 Humerus_ectepicondylar_groove /  present absent,
106 Humerus_ectepicondyle /  absent present,
107 Humerus_entepicondyle /  absent present,
108 Radius_shape /  not_expanded slightly_expanded_anterodistally broadly_expanded,
109 Ulna_contact_with_centrale /  absent present,
110 Radiale_size /  'large, broad' small_to_absent,
111 Carpal_reduction /  six_or_more_carpals five_or_less_carpals,
112 Pisiform /  present absent,
113 Metacarpal_I_expansion /  'elongate, spindle-shaped' broadly_expanded,

```
          114 Phalanx_shape /  'elongate, spindle-shaped' 'blocky,
hourglass-shaped',
          115 Ilium_crest /  'blade-like, articulates with sacral ribs'
'elongate, cylindrical, does not articulate with sacral ribs',
          116 Ilium_acetabular_area /
arcuate_ridge_supertending_acetabulum 'acetabulum set into broad, short
V-shaped notch',
          117 Pubic_tubercle_condition /
tubercle_an_elongate_protuberance_near_midpoint_of_shaft 'thin,
semicircular crest-like blade close to the acetabulum' absent,
          118 Ischiadic_tubercle_size /  elongate short,
          119 Astragalus /  notched_emargination_for_the_crural_foramen
'without notch, pedunculate fibular articulation present',
          120 Appendicular_epiphyses /  formed_from_ossified_cartilage
from_thick_unossified_cartilage epiphyses_missing_or_extremely_thin,
          121 Hyperphalangy /  absent present,
          122 Posterior_thoracic_vertebra /
not_markedly_longer_than_anterior_dorsal_vertebrae
markedly_longer_than_anterior_dorsal_vertebrae,
          123 Ecp_process_of_pterygoid /  'distal portion not offset
anterolaterally, &/or lacking longitudinal grooves & ridges' 'distal
portion of process offset anterolaterally, bears longitudinal grooves &
ridges',
          124 'Quadrate mid-shaft lateral deflection' /  absent
present,
          125 'Caudal vertebrae, depth' /  'no sculling organ, ratio
being 1,4 or less' 'weakly developed sculling organ, ratios varying from
1,4 to 2,5' 'very deep, forming a strong sculling organ, ratio being 2.6
or higher',
          126
Posteromedial_canal_enters_basisphenoid_between_the_carotid_artery_path_a
nd_abducens_nerve_exit /  absent present,
          127
Vascularization_of_the_basisphenoid_and_basioccipital_type /  _type_1_
type_2 type_3,
          128
_Dorsal_surface_of_the_frontal_bears_discrete_ornamentations /  absent
present,
          129 Retroarticular_process_lenght /  elongated short
extremely_short ;
      MATRIX
      'Shinisaurus_crocodilurus'    001000000001003-
10010110001030201000000-0001000000{2 3}0---1000000101000010000-
000000001000000100-0?000000000000000000000010000{0 1}000000
      'Lanthanotus_borneensis'      001000100011003-00---
1000?103022001000000-001100000051---1000000100001010100-
000000001200000010-0?000000000000000000000110000{0 1}000000
      'Varanus_komodoensis'         000000000001003-100101000?10302000--
0100-0011000000{4 5}1---1000000100000011000-000000001100{0 1}00000-
0?000000000000000000000010000{0 1}000000
      'Heloderma_horridum'          000010000011003-00---
1100?10302010000000-000100000050---1000000100000010000-000000001200{0
1}00100-0?010000000000000000000010000{0 1}000000
```

'Estesia_mongoliensis' 000000000001?03-00---
1000?1030?100000?00?0001????0051---
10?00???0000010?????????00?0?????????????????00000?00??????0?????0??{0
1}0?0000
'Adriosaurus_suessi'
000???10101??000??10?0??100???1??0?????????0?????????????????0?0???????????
?1?1?0???01201000??????010??????1????1100???100??1??00
'Dolichosaurus_longicollis'
???????????????????????????????????????????????????????00??????????????????
?1011000000130??00?10-?0001???????1??????0????2?0??1????
'Pontosaurus_kornhuberi' 00010101101??03-
0000?11?0010?0010???0??0?000?0?0????0???????0??00?0100??1010100???012000
00000-012?0????111?00?01001?1000?01??00
'Aigialosaurus_bucchichi' 00??1?12?00??03-
0000?1101010??0100001?00?0110000100300?00?00110??000000-
1???0?00????00?0000?01000000????000000000????1?00002??01
'Aigialosaurus_dalmaticus' ????0?12100??03-
0000?1101010??110???3????011?0???00????00???02001?0000??10???000???0000??
??????????0????00000000000001?1000?02??01
'Komensaurus_carrolli'
?????????????????????????????????????20?0?010?0?????????????????0000?????????
?1???0?01??0?0?00??????????????????0000?000?0011?000?02????
Portunatasaurus_krambergeri
???????????????????????????????????????????????????????????????????????????
???1?00????00??????10-0?000??00111110001 0????001???????
'Halisaurus_platyspondylus' 010000021100003-
1010100000???10???102002001 0?0001?????101?0001000100001 0??0?????00???????
1???????????????????????????1?????00020001
'Eonatator_sternbergii' 0100?0021000003-
0010100000??020???0020?2?010?0?01?0?01101?0?0100010?0010?10??00100??01110
101111100100100001011100111 1?010?020001
'Pluridens_serpentis'
?01?0?00110?????1011?1?0011?12021???2???2??10?0?????010??1???1?000?{0
1}?{0 1}010??????????????????????????????????????????????????????0001
'Yaguarasaurus_columbianus'
010?0?02000??1201200?110000?302111??2100200110000013 11???????????0000010?
?1??????????????????????????????????????????????0?0?120?
'Romeosaurus_fumanensis'
??????02??????????????????1???20?21?102100201 1?2011??211??00000100?1000010?
????01101??0?????????0-11??00?00?00???????????????????10???01
'Russellosaurus_coheni'
11??0002000111201200011000012022110021002011120000120110000000100100000 10?
??????????????????????????????????????????????10?1101
Garamaudo_beauciensis
?????????????????????????????????????????????????????????0?01000????????
?1??0?1?11????????????????????00000?11????????00?????????????2
'Mosasauroid_of_Villeveyrac'
??????02?101013?????????????0??????????????????????????????????????????????
?1??0010?0???????0?????????????????????????0????????????????1?
'Pannoniasaurus_inexpectatus'
00000???????????????????????01?????????1100201 0?0000??0?20010?0010000000?11?
?10?001000???0??0?????????????0001011????????00????????0?????2

```
      'Tethysaurus_nopcsai'
000?0?021100?1201200?110000?10?21?0001002010 1?20001101100100010000 00{0
1}00-1010?001000??????0??10-0000????011???????000?????00?1112
      'Taniwhasaurus_antarcticus'   31111?00?210?23-
10010110020030?21???11?02000?0010???231000?00?0??0111011?0??????1?1?????0
0??????????????????????????????0?1?000?
      'Taniwhasaurus_oweni'
31111?00?21??2????01????020030?21?0011?02000?0010??323??00?0?????011101?1
00????01010????00??1100101???????????????????????0?0?000?
      'Tylosaurus_nepaeolicus'      31111?001{0 1}10123-
1001?1100200301211001100211110010 0?42211000000000000010100??00010101???0
0??????????0001000101110 0???????00020001
      'Tylosaurus_pembinensis'      31?1??001?10123-
1001?1100200301 2??0011101210?00100?{4
5}2210000000000101010111 00??000111011?0?01011011011000100 01??????0110?1100
0?0001
      'Tylosaurus_proriger'         311111001110?23-1001011002003{0
1}12110011101000100100042211000000000000{0 1}001{0
1}1000000010101010001011011011000100010111001020011000 20001
      'Angolasaurus_bocagei'
11??0?01?11??1??1100?1??000?30???1??2100010010001015021?010001001?00101??
?1????????????????????????????????????????????0?0?1201
      'Ectenosaurus_clidastoides'
2100010200111120111000 1?0001112110001001210111 0?1021120001000000100010100
01001001???0???????01011 0021???00120101 00?????11100?1201
      'Gavialimimus_ptychodon'      010?0?10001?1?0-
121111??0110310???00?1??1?0????????503????????11??1001010??0??1?0?????????
??????????2?????11??????????????????0?????0?
      'Goronyosaurus_nigeriensis'
01000?10?111????1?10??????????????00????????????????5021????????????????01??
??????????????????????????????????????????????????0?????0?
      'Latoplatecarpus_nichollsae'  01000?10?210?10-
1112?1??010?300???0021?00200?0011025011???????????001101?1011???0?????????
??????????21111?????????????????????0?120?
      'Latoplatecarpus_willistoni'  01000?0012011?0-
11120?1?010130121?00210012001011102502110100000001000101011100010 11??0????
0???2111??????????????????????????00?1201
      'Platecarpus_tympaniticus'    010000001111110-
1111011001013011 1?002110110010111025021101000000010000010100-
000011000010001001011002 0??1?01201110010110110 0021201
      'Plesioplatecarpus_planifrons'110000001011?10-
1110?110000130 1?100021001000?0011015121001000000010001010101000 0011??0????
???01011002 0?1100?2011100?????1?000?1201
      'Plioplatecarpus_primaevus'   010???001210010-
1112011?010030010000 21100200?0111015020101000000010000010100-
000011000 11??0??010110020111001201110010 20?1?000?1201
      'Selmasaurus_johnsoni'
11000?1000101120120001 1?0001?0?21?0011011100?000101502111100000001000 10101
010000011000??????????????????????????????????????1?1201
      'Selmasaurus_russelli'        ??????100101?0-
121010????01???21???2111100?100010 2??????????????????????10???????????????
??????????????????????????????????1?120?
```

'Dallasaurus_turneri'
??????0??0?????0???????????????????0?????????????????1010??0?1????10001??
11??00111110????1???0-0???01000000????????0???0?1?????1?
'Clidastes_liodontus'
21000?02001001101001111?10102112011010?010000001010211101000100001 0{0
1}00100110000011010311110101100002101011121000101000121 0?020001
'Clidastes_moorevillensis'
2100000200100110100111 1?1010211 20?1010?000100001010212101000100001010010 0
110000011?1031111??0110000210101112100010100012 11???0001
'Clidastes_propython'
21000?1200100110100111??1010211???1010?0001000010101121 01?101000010100100
110000011?10??1?1??????0002101011121 0?0???????2?10020001
'Eremiasaurus_heterodontus'
11001?10111??11?10?1?11?1?0130120?111??1??0??1?111?2121010??1011?10001110
1??0100110?0?1111001?01??10?????????????01??01??1??20001
'Gnathomortis_stadtmani'
11000????10??0?1001?111??1?30????1010?00110?0011??3?2??10?01?01?10{0
1}?010011????0???1?????1???????????????????????????????????0001
'Mosasaurus_conodon'
?????????????????????????????????????????????????????21010???000010100110
11100101101 1????1???1?00103111011 1?1??????????2?1?0?00?1
'Mosasaurus_hoffmanni'
210210000110011110021 11?110031221010000010 1?11011031210101110110101001 10
1110110110?1?11111?02100113111011121000111000121 10020001
'Mosasaurus_missouriensis'
21000?10011??1111001?11?1?00302???1000?00100?1?011?312?01011101101010 01?0
11?00001101131??11?01??00?????????2100?11?????2?1?020001
'Plesiotylosaurus_crassidens'
3101??10?11??111100111 1?120031 2??01100?10110?0?101?212111011100??1010011?
????00011?10??????0110011312101112100010?????2???0?000?
'Plotosaurus_bennisoni'
11001?101010?1111011?1111?001112001000?001000110 11?1211?1???10110000011?0
10?10?011??13??1?1?021001131110111210001 1??1??211?020001
'Prognathodon_currii'
100???0111????10??11?1?0030220??1???????0????1??50??????????000??100?10?
????????????????????????????????????????????????????0001
'Prognathodon_kianda'
11001??00?1???1?10?1?11??0??30?2111?10?10110000101?2121?1000100001000110?
?????????????????????????????????????????????????00?0001
'Prognathodon_overtoni'
11000?10011001111001?1111?10302200111011011 0?001010312101011100101100111?
???0?0?110?13?11111011100121210111??0???010000211?020001
'Prognathodon_rapax'
11000?10001001111001111?1?1?31120?1110?1011000011 10?021010???000011001110
110?00011110????1??01?00112121011121 00010?????2?1???0001
'Prognathodon_saturator'
??00??10?110011110111???1000???20?111?010110000101?3?2101??0100101100 1110
?10?01011110??01??0???????????????2?????????????????0001
'Prognathodon_solvayi'
000000100010011010111111100030210?1110010100?001111402?010101?00000?01110
?10001011?10???????01?01??????????????????????????0?0001
'Prognathodon_waiparaensis'
110000?????????????????????????21?111??10110?0010??3??101????????00?0111?
?????????????????????????????????????????????????0??00?1

```
      'Globidens_alabamaensis'
??????10?111011?????????1010?1????1010?111?0??010?????1010?11000011001100
11??0?011??0???????????0??21010111????????????2?1???000?
      'Globidens_dakotensis'
110?0?10111??1101001?11?1?1031120?1010?10110?0?1010??????????????11001100
11?00?011?10????????????????????????????????????10??000?
      'Globidens_schurmanni'
011????0?0??????101??????21?31??????1??11?10???11???1?111?1???????10010-
01???010111??????1??020?10121?????1?100?10?????????0?000?
      'Jormungandr_walhallaensis'
3102100101100110????????101121?10?1010001110?1011??22211100010000????01?0
1110010??????????????????????????????????????????00?0001
      Vallecillosaurus_donrobertoi
?????????????????????????????????????????????????????????????????????????
?????????????01100??????????????????????01?0?000???2????
      Haasiasaurus_gittelmani
????????????????????????????????????11?02?00???????00010100000000000001??
?1000000100000?0???11100100?000000000000000???000??????1
      Carsosaurus_marchesetti
?????????????????????????????????????????????????????????????????????????
?1???00????000?10??10-0001000000100000000001??001??2????

;

END;
BEGIN ASSUMPTIONS;
      TYPESET * UNTITLED   =  unord:  1- 7 9- 18 21- 22 24- 26 28 30- 31
33- 35 38- 40 42- 49 51 53- 59 61- 74 77- 78 80- 81 84 88 90- 91 95- 99
101- 110 112- 117 119- 124 126 128- 129, ord:  8 19- 20 23 27 29-32\3 36-
37 41 50 52 60 75- 76 79-82\3 83 85- 87 89-92\3 93- 94 100 111 118 125
127;

END;

BEGIN MESQUITECHARMODELS;
      ProbModelSet * UNTITLED   =  'Mk1 (est.)':  1- 129;
END;

BEGIN NOTES;

      TEXT   TAXON = 28 CHARACTER = 126 TEXT = 1;

      TEXT   CHARACTER = 26 STATE = 0 TEXT = 'absent^n';
      TEXT   CHARACTER = 47 STATE = 0 TEXT = single_thin;

            MESQUITESCRIPTVERSION 2;
            TITLE AUTO;
            tell ProjectCoordinator;
            timeSaved 1676237305457;
            getEmployee #mesquite.minimal.ManageTaxa.ManageTaxa;
            tell It;
                  setID 0 2466279084245358266;
            endTell;
```

```
			getEmployee
#mesquite.charMatrices.ManageCharacters.ManageCharacters;
			tell It;
				setID 0 2717975509959130391;
				mqVersion 370;
				checksumv 0 3 3313146721 null getNumChars 124 numChars
124 getNumTaxa 58 numTaxa 58 short true bits 2305843009213694015 states
63 sumSquaresStatesOnly 36352.0 sumSquares -7.378697629483819E19
longCompressibleToShort false usingShortMatrix true NumFiles 1
NumMatrices 1;
				mqVersion;
			endTell;
			getWindow;
			tell It;
				suppress;
				setResourcesState false false 122;
				setPopoutState 300;
				setExplanationSize 0;
				setAnnotationSize 0;
				setFontIncAnnot 0;
				setFontIncExp 0;
				setSize 805 636;
				setLocation 8 25;
				setFont SanSerif;
				setFontSize 10;
				getToolPalette;
				tell It;
				endTell;
				desuppress;
			endTell;
			getEmployee
#mesquite.charMatrices.BasicDataWindowCoord.BasicDataWindowCoord;
			tell It;
				showDataWindow #2717975509959130391
#mesquite.charMatrices.BasicDataWindowMaker.BasicDataWindowMaker;
				tell It;
					getWindow;
					tell It;
						setExplanationSize 30;
						setAnnotationSize 20;
						setFontIncAnnot 0;
						setFontIncExp 0;
						setSize 683 564;
						setLocation 8 25;
						setFont SanSerif;
						setFontSize 10;
						getToolPalette;
						tell It;
							setTool
mesquite.charMatrices.BasicDataWindowMaker.BasicDataWindow.ibeam;
						endTell;
						setTool
mesquite.charMatrices.BasicDataWindowMaker.BasicDataWindow.ibeam;
```

```
				colorCells
#mesquite.charMatrices.NoColor.NoColor;
				colorRowNames
#mesquite.charMatrices.TaxonGroupColor.TaxonGroupColor;
				colorColumnNames
#mesquite.charMatrices.CharGroupColor.CharGroupColor;
				colorText
#mesquite.charMatrices.NoColor.NoColor;
				setBackground White;
				toggleShowNames on;
				toggleShowTaxonNames on;
				toggleTight off;
				toggleThinRows off;
				toggleShowChanges on;
				toggleSeparateLines off;
				toggleShowStates on;
				toggleReduceCellBorders off;
				toggleAutoWCharNames on;
				toggleAutoTaxonNames off;
				toggleShowDefaultCharNames off;
				toggleConstrainCW on;
				toggleBirdsEye off;
				toggleColorOnlyTaxonNames off;
				toggleShowPaleGrid off;
				toggleShowPaleCellColors off;
				toggleShowPaleExcluded off;
				togglePaleInapplicable on;
				togglePaleMissing off;
				toggleShowBoldCellText off;
				toggleAllowAutosize on;
				toggleColorsPanel off;
				toggleDiagonal on;
				setDiagonalHeight 80;
				toggleLinkedScrolling on;
				toggleScrollLinkedTables off;
			endTell;
			showWindow;
			getWindow;
			tell It;
				forceAutosize;
			endTell;
			getEmployee
#mesquite.charMatrices.AlterData.AlterData;
			tell It;
				toggleBySubmenus off;
			endTell;
			getEmployee
#mesquite.charMatrices.ColorByState.ColorByState;
			tell It;
				setStateLimit 9;
				toggleUniformMaximum on;
			endTell;
			getEmployee
#mesquite.charMatrices.ColorCells.ColorCells;
```

```
                    tell It;
                          setColor Red;
                          removeColor off;
                    endTell;
                    getEmployee
#mesquite.categ.StateNamesEditor.StateNamesEditor;
                    tell It;
                          makeWindow;
                          tell It;
                                setExplanationSize 30;
                                setAnnotationSize 20;
                                setFontIncAnnot 0;
                                setFontIncExp 0;
                                setSize 683 564;
                                setLocation 8 25;
                                setFont SanSerif;
                                setFontSize 10;
                                getToolPalette;
                                tell It;
                                      setTool
mesquite.categ.StateNamesEditor.StateNamesWindow.ibeam;
                                endTell;
                                setActive;
                                rowsAreCharacters on;
                                toggleConstrainChar on;
                                toggleConstrainCharNum 3;
                                togglePanel off;
                                toggleSummaryPanel off;
                          endTell;
                          showWindow;
                    endTell;
                    getEmployee
#mesquite.categ.StateNamesStrip.StateNamesStrip;
                    tell It;
                          showStrip off;
                    endTell;
                    getEmployee
#mesquite.charMatrices.AnnotPanel.AnnotPanel;
                    tell It;
                          togglePanel off;
                    endTell;
                    getEmployee
#mesquite.charMatrices.CharReferenceStrip.CharReferenceStrip;
                    tell It;
                          showStrip off;
                    endTell;
                    getEmployee
#mesquite.charMatrices.QuickKeySelector.QuickKeySelector;
                    tell It;
                          autotabOff;
                    endTell;
                    getEmployee
#mesquite.charMatrices.SelSummaryStrip.SelSummaryStrip;
                    tell It;
```

```
                                showStrip off;
                          endTell;
                          getEmployee
#mesquite.categ.SmallStateNamesEditor.SmallStateNamesEditor;
                          tell It;
                                panelOpen true;
                          endTell;
                    endTell;
              endTell;
              endTell;

END;

Begin MESQUITE;
              MESQUITESCRIPTVERSION 2;
              TITLE AUTO;
              tell ProjectCoordinator;
              timeSaved 1765819813191;
              getEmployee #mesquite.minimal.ManageTaxa.ManageTaxa;
              tell It;
                    setID 0 5742113724525426697;
                    tell It;
                          setDefaultOrder  0 1 2 3 4 5 6 10 7 8 9 59 11 12
13 14 15 16 57 58 17 18 19 20 21 22 23 24 25 26 27 28 29 30 31 32 33 34
35 36 37 38 39 40 41 42 43 44 45 46 47 48 49 50 51 52 53 54 55 56 60 61
62;
                          attachments ;
                    endTell;
              endTell;
              getEmployee
#mesquite.charMatrices.ManageCharacters.ManageCharacters;
              tell It;
                    setID 0 8407246798674527177;
                    tell It;
                          setDefaultOrder  0 1 2 3 4 5 6 7 8 9 10 11 12 13
14 15 16 17 18 19 20 21 22 23 24 25 26 27 28 29 30 31 32 33 34 35 36 37
38 39 40 41 42 43 44 45 46 47 48 49 50 51 52 53 54 55 56 57 58 59 60 61
62 63 64 65 66 67 68 69 70 71 72 73 74 75 76 77 78 79 80 81 82 83 84 85
86 87 88 89 90 91 92 93 94 95 96 97 98 99 100 101 102 103 104 105 106 107
108 109 110 111 112 113 114 115 116 117 118 119 120 121 122 123 124 125
128 126 127;
                          attachments ;
                    endTell;
                    mqVersion 381;
                    checksumv 0 3 548116782 null  getNumChars 129 numChars
129 getNumTaxa 63 numTaxa 63   short true   bits 2305843009213694015
states 63   sumSquaresStatesOnly 37412.0 sumSquares -7.378697629483819E19
longCompressibleToShort false usingShortMatrix true   NumFiles 1
NumMatrices 1;
                    mqVersion;
              endTell;
              getWindow;
              tell It;
                    suppress;
```

```
                  setResourcesState false false 100;
                  setPopoutState 300;
                  setExplanationSize 0;
                  setAnnotationSize 0;
                  setFontIncAnnot 0;
                  setFontIncExp 0;
                  setSize 1707 833;
                  setLocation -8 0;
                  setFont SanSerif;
                  setFontSize 10;
                  getToolPalette;
                  tell It;
                  endTell;
                  desuppress;
            endTell;
            getEmployee
#mesquite.charMatrices.BasicDataWindowCoord.BasicDataWindowCoord;
            tell It;
                  showDataWindow #8407246798674527177
#mesquite.charMatrices.BasicDataWindowMaker.BasicDataWindowMaker;
                  tell It;
                        getWindow;
                        tell It;
                              getTable;
                              tell It;
                                    rowNamesWidth 186;
                              endTell;
                              setExplanationSize 30;
                              setAnnotationSize 20;
                              setFontIncAnnot 0;
                              setFontIncExp 0;
                              setSize 1607 761;
                              setLocation -8 0;
                              setFont SanSerif;
                              setFontSize 14;
                              getToolPalette;
                              tell It;
                                    setTool
mesquite.charMatrices.BasicDataWindowMaker.BasicDataWindow.ibeam;
                              endTell;
                              setActive;
                              setTool
mesquite.charMatrices.BasicDataWindowMaker.BasicDataWindow.ibeam;
                              colorCells
#mesquite.charMatrices.NoColor.NoColor;
                              colorRowNames
#mesquite.charMatrices.TaxonGroupColor.TaxonGroupColor;
                              colorColumnNames
#mesquite.charMatrices.CharGroupColor.CharGroupColor;
                              colorText
#mesquite.charMatrices.NoColor.NoColor;
                              setBackground White;
                              toggleShowNames on;
                              toggleShowTaxonNames on;
```

```
					toggleTight off;
					toggleThinRows off;
					toggleShowChanges on;
					toggleSeparateLines off;
					toggleShowStates on;
					toggleReduceCellBorders off;
					toggleAutoWCharNames on;
					toggleAutoTaxonNames off;
					toggleShowDefaultCharNames off;
					toggleConstrainCW on;
					toggleBirdsEye off;
					toggleColorOnlyTaxonNames off;
					toggleShowPaleGrid off;
					toggleShowPaleCellColors off;
					toggleShowPaleExcluded off;
					togglePaleInapplicable on;
					togglePaleMissing off;
					toggleShowBoldCellText off;
					toggleAllowAutosize on;
					toggleColorsPanel off;
					toggleDiagonal on;
					setDiagonalHeight 80;
					toggleLinkedScrolling on;
					toggleScrollLinkedTables off;
				endTell;
				showWindow;
				getWindow;
				tell It;
					forceAutosize;
				endTell;
				getEmployee
#mesquite.charMatrices.AlterData.AlterData;
				tell It;
					toggleBySubmenus off;
				endTell;
				getEmployee
#mesquite.charMatrices.ColorByState.ColorByState;
				tell It;
					setStateLimit 9;
					toggleUniformMaximum on;
				endTell;
				getEmployee
#mesquite.charMatrices.ColorCells.ColorCells;
				tell It;
					setColor Red;
					removeColor off;
				endTell;
				getEmployee
#mesquite.categ.StateNamesEditor.StateNamesEditor;
				tell It;
					makeWindow;
					tell It;
						getTable;
						tell It;
```

```
                                   rowNamesWidth 581;
                              endTell;
                              setExplanationSize 30;
                              setAnnotationSize 20;
                              setFontIncAnnot 0;
                              setFontIncExp 0;
                              setSize 1607 761;
                              setLocation -8 0;
                              setFont SanSerif;
                              setFontSize 14;
                              getToolPalette;
                              tell It;
                                   setTool
mesquite.categ.StateNamesEditor.StateNamesWindow.ibeam;
                              endTell;
                              rowsAreCharacters on;
                              toggleConstrainChar on;
                              toggleConstrainCharNum 3;
                              togglePanel off;
                              toggleSummaryPanel off;
                         endTell;
                         showWindow;
                    endTell;
                    getEmployee
#mesquite.categ.StateNamesStrip.StateNamesStrip;
                    tell It;
                         showStrip off;
                    endTell;
                    getEmployee
#mesquite.charMatrices.AnnotPanel.AnnotPanel;
                    tell It;
                         togglePanel off;
                    endTell;
                    getEmployee
#mesquite.charMatrices.CharReferenceStrip.CharReferenceStrip;
                    tell It;
                         showStrip off;
                    endTell;
                    getEmployee
#mesquite.charMatrices.QuickKeySelector.QuickKeySelector;
                    tell It;
                         autotabOff;
                    endTell;
                    getEmployee
#mesquite.charMatrices.SelSummaryStrip.SelSummaryStrip;
                    tell It;
                         showStrip off;
                    endTell;
                    getEmployee
#mesquite.categ.SmallStateNamesEditor.SmallStateNamesEditor;
                    tell It;
                         panelOpen true;
                    endTell;
               endTell;
```

```
            endTell;
            endTell;
end;
```
